# Proteome capacity constraints favor respiratory ATP generation

**DOI:** 10.1101/2022.08.10.503479

**Authors:** Yihui Shen, Hoang V. Dinh, Edward Cruz, Catherine M. Call, Heide Baron, Rolf-Peter Ryseck, Jimmy Pratas, Arjuna Subramanian, Zia Fatma, Daniel Weilandt, Sudharsan Dwaraknath, Tianxia Xiao, John I. Hendry, Vinh Tran, Lifeng Yang, Yasuo Yoshikuni, Huimin Zhao, Costas D. Maranas, Martin Wühr, Joshua D. Rabinowitz

**Affiliations:** Department of Chemistry, Princeton University, Princeton, NJ, USA; Lewis Sigler Institute for Integrative Genomics, Princeton University, Princeton, NJ, USA; Ludwig Institute for Cancer Research, Princeton Branch, Princeton, NJ, USA; Department of Molecular Biology, Princeton University, Princeton, New Jersey 08544, USA; Department of Chemical Engineering, The Pennsylvania State University, University Park, PA, USA; Carl R. Woese Institute for Genomic Biology, University of Illinois Urbana-Champaign, Urbana, IL, 61801, USA; Department of Chemical and Biomolecular Engineering, University of Illinois at Urbana-Champagne, Urbana, IL, USA; US Department of Energy Joint Genome Institute, Lawrence Berkeley National Laboratory, Berkeley, CA 94720, USA

## Abstract

Cells face competing metabolic demands. These include efficient use of both limited substrates and limited proteome capacity, as well as flexibility to deal with different environments. Flexibility requires spare enzyme capacity, which is proteome inefficient. ATP generation can occur via fermentation or respiration. Fermentation is much less substrate-efficient, but often assumed to be more proteome efficient ^1–3^, thereby favoring fast-growing cells engaging in aerobic glycolysis ^4–8^. Here, however, we show that mitochondrial respiration is actually more proteome-efficient than aerobic glycolysis. Instead, aerobic glycolysis arises from cells maintaining the flexibility to grow also anaerobically. These conclusions emerged from an unbiased assessment of metabolic regulatory mechanisms, integrating quantitative metabolomics, proteomics, and fluxomics, of two budding yeasts, *Saccharomyces cerevisiae* and *Issatchenkia orientalis*, the former more fermentative and the latter respiratory. Their energy pathway usage is largely explained by differences in proteome allocation. Each organism’s proteome allocation is remarkably stable across environmental conditions, with metabolic fluxes predominantly regulated at the level of metabolite concentrations. This leaves extensive spare biosynthetic capacity during slow growth and spare capacity of their preferred bioenergetic machinery when it is not essential. The greater proteome-efficiency of respiration is also observed in mammals, with aerobic glycolysis occurring in yeast or mammalian cells that maintain a fermentation-capable proteome conducive to both aerobic and anaerobic growth.

## Introduction

Metabolism is subject to physical constraints. Given the law of conservation of matter, to maintain homeostasis, limited metabolic inputs must balance with outputs (‘flux balance’). These outputs include high energy cofactors (most importantly ATP), building blocks for cell replication, and waste. The resources, including physical space and protein synthesis capacity, to sustain these fluxes are also limited. Thus, cells are under pressure to produce their required metabolic fluxes efficiently. As proteins catalyze most metabolic reactions and comprise the majority of biomass in many cell types, the challenges of limited biosynthetic machinery and physical space can be viewed largely as constraints on proteome capacity ^2,3,9–17^.

To manage proteome capacity, cells tailor protein expression to their conditions ^18–21^. For example, rapidly growing cells express copious ribosomes ^18,22,23^. This requires less expression of other protein types, such as anabolic and catabolic enzymes (e.g. those required for assimilating limiting quantities of nitrogen ^24^, or breaking down non-preferred carbon sources ^21,25^). Conversely, under less favorable conditions, ribosome expression falls to make room for other proteins. Perfect proteome tailoring, however, is not necessarily feasible or desirable ^26–30^, as proteome remodeling is expensive and spare enzyme capacity can allow cells to quickly ramp up fluxes to deal with changing conditions.

An important case of metabolic tailoring involves energy production from fermentation versus respiration. Respiration is by far more energy-efficient, producing roughly 10-fold more ATP per glucose ^31^. Nevertheless, many organisms ferment, producing organic waste, even when oxygen is available (‘aerobic glycolysis’). Aerobic glycolysis is associated with fast growing cells including bacteria, yeast and cancer cells ^4–8^. Indeed, as their growth accelerates, both *Escherichia coli* and *S. cerevisiae* switch from respiration to fermentation ^2,32^.

Why do cells engage in aerobic glycolysis when it is so much less energy-efficient than respiration? One possibility involves a ‘rate-yield tradeoff ^1–3,33^. More specifically, it is often believed that fermentation is capable of producing ATP faster per unit enzyme expression, i.e. more ‘proteome efficient’^9–17^. The proteome efficiency of glycolysis versus respiration, however, has not been carefully experimentally tested in eukaryotes.

Here we examine this question, building from an extensive systems-level analysis of metabolism in two evolutionarily distant budding yeasts (separated by 200 million years ^34^): *S. cerevisiae* (Baker’s yeast) and *I. orientalis* (also known as *Candida krusei* and *Pichia kudriavzevii*, a species abundant in fermented food, fruit, and soil with favorable properties for bioengineering) ^35–40^. We find that, across diverse environmental conditions spanning a broad range of growth rates, the proteome of each yeast varies only modestly, with metabolic flux explained primarily by metabolite levels. Across the yeasts, however, metabolic flux differences are explained mainly by proteome allocation, with *I. orientalis* expressing more respiratory enzymes and outcompeting *S. cerevisiae* across diverse aerobic contexts. Both in *I. orientalis* and in respiring *S. cerevisiae*,quantitative measurements show that respiration is actually several-fold more proteome efficient than glycolysis. Similar results are attained in mammalian tissues and cancer cells. The origin of aerobic glycolysis accordingly does not lie in proteome efficiency, but rather in a proteome hedging strategy where cells maintain spare glycolytic capacity in preparation for potential future hypoxia.

## Results

### Metabolic fluxes in *S. cerevisiae* and *I. orientalis*

We first characterized aerobic growth and metabolism of *S. cerevisiae* and *I. orientalis* in glucose minimal medium (Fig. S1a). *I. orientalis* grows faster than *S. cerevisiae* (μ = 0.52 vs. 0.39 h^-1^), respires much more, consumes less glucose, and excretes less ethanol (Fig. S1a). We then resolved flux through the entire metabolic network with ^13^C metabolic flux analysis (Fig. 1). Specifically, we developed genome-scale metabolic models of both yeasts including complete atom mapping ^41^. The models were then constrained by flux balance and experimentally derived extracellular fluxes, biomass fluxes, and isotope labeling (from two distinct ^13^C-tracer strategies: [1,2-^13^C_2_]glucose and [U-^13^C_6_]glucose, each at 1:1 molar ratio with unlabeled glucose). This enabled comprehensive yeast metabolic flux analysis, at a rigor previously achieved only in prokaryotes ^41,42^.

**Figure 1.**
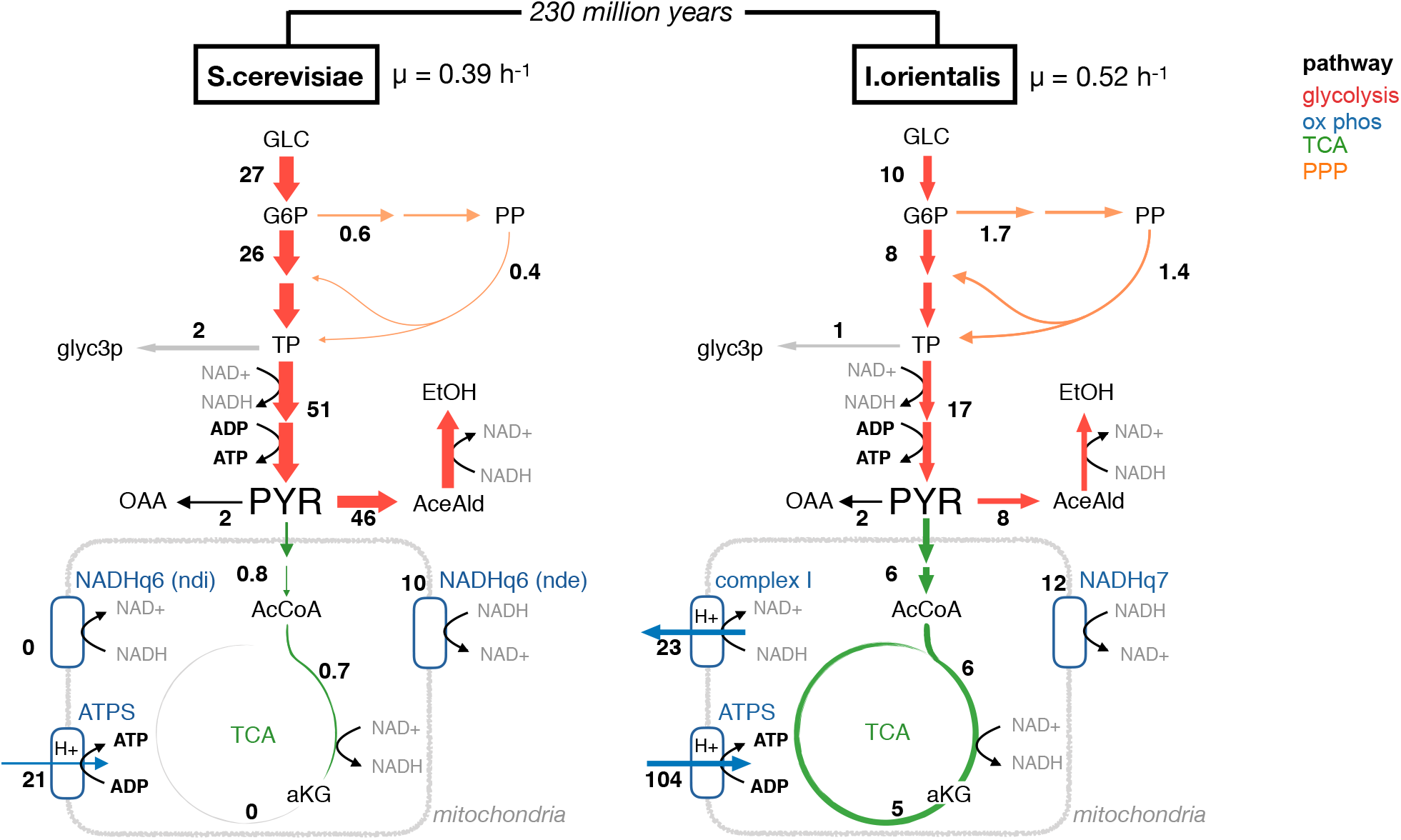
Genome-scale flux analysis shows more active respiratory metabolism in *I. orientalis*. Metabolic flux (in mmol/gDW/h) of *S. cerevisiae* (CEN.PK) and *I. orientalis* (SD108) in aerobic exponential growth in YNB with 20 g/L glucose. Fluxes are best estimate from genome-scale ^13^C-informed metabolic flux analysis (MFA), with the input data including metabolite ^13^C labeling from two ^13^C-glucose tracers (each with n = 3 or 4 biological replicates) and consumption and excretion fluxes (at least n = 3 biological replicates). Color represents metabolic pathways: glycolysis in red, oxidative phosphorylation (ox phos) in blue, TCA in green, and PPP in orange. Numbers represents flux in mmol/h/gDW.

^13^C-tracing resolved key internal flux branch points. For example, [1,2-^13^C_2_]glucose revealed markedly higher [M+1] pyruvate labeling in *I. orientalis* (Fig. S1b), reflecting greater oxidative pentose phosphate pathway (PPP) flux in this yeast species (Fig. 1). The same tracer also reveals greater [M+1] tricarboxylic acid (TCA) cycle intermediates in *I. orientalis* (Fig. S1c), consistent with higher oxidative TCA cycle flux relative to *S. cerevisiae*, where clockwise TCA flux was truncated at α-ketoglutarate (Fig. 1). The greatest difference between the two yeasts is in the way glucose is catabolized. Namely, *S. cerevisiae* prefers carbon-inefficient fermentation, and correspondingly makes most ATP from glycolysis and consumes most NADH via ethanol fermentation (Fig. 1, Fig. S1d). In contrast, *I. orientalis* prefers respiration for ATP production, and re-oxidizes NADH by a blend of complex I and the quinone oxidoreductase Nde1, the knockout of which impairs *I. orientalis* but not *S. cerevisiae* growth ^43^ (Fig. 1, Fig. S1e).

### Flux control across environmental conditions

When nutrients become scarce, cells adjust metabolic fluxes and growth. Such fluxes can be controlled through enzyme concentration (k_cat_[E] in the Michaelis-Menten kinetics), active site occupancy ([S]/([S]+K_m_)), or allosteric regulation (Fig. 2a). To assess flux control mechanisms in *S. cerevisiae* and *I. orientalis*, we grew each yeast in aerobic chemostats at diverse growth rates controlled by limiting glucose, ammonia, or phosphate availability (Fig. 2b). In each nutrient environment, we measured enzyme concentrations via quantitative proteomics, metabolite concentrations via metabolomics, and metabolic fluxes via ^13^C-informed fluxomics (Fig. 2c). Fluxes aligned remarkably closely with growth rate in both yeasts (Fig. S2a), with the exception of metabolic switching to respiration and the PPP (relative to glycolysis) in glucose-limited *S. cerevisiae*, which renders glucose-limited *S. cerevisiae* metabolically similar to *I. orientalis* (Fig. S2, b and c). On average, 53% of flux variation in *S. cerevisiae* and 71% in *I. orientalis* was explained by growth rate alone.

**Figure 2.**
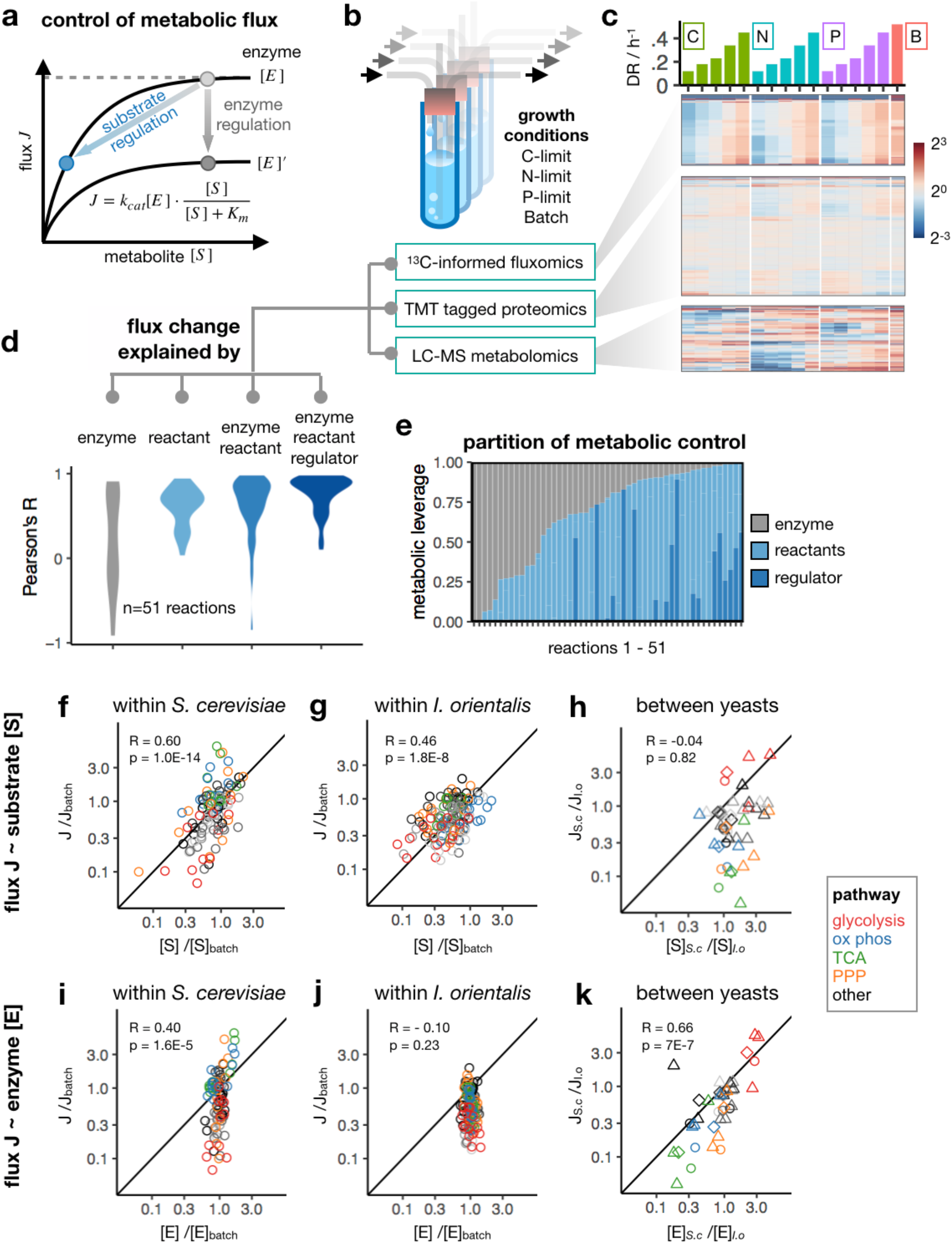
Flux change is explained by metabolites within yeast species, and by enzymes across yeasts. (a) Flux change can be achieved through change in either enzyme or substrate level via Michaelis-Menten kinetics. (b) Multi-omics data were obtained in steady-state yeast grown in nutrient-limited continuous culture or nutrient-replete batch culture. (c) Genome-wide metabolic flux, protein abundance, and metabolite concentrations in *I. orientalis* across growth conditions. Values are normalized to the geometric mean across all the conditions. Metabolomics, mean of n = 3 technical replicates (independent sampling from continuous culture). Proteomics, n = 1 biological replicate. Fluxomics, best estimates from ^13^C-MFA similar to Fig. 1. (d) Distribution of Pearson’s R for 51 reactions between measured flux and flux predicted from Michaelis-Menten kinetics accounting for different variables: concentration of enzyme, reactant, and best data-supported allosteric regulator (if any). (e) Partition of metabolic control among enzyme, reactants, and regulator for 51 reactions in *I. orientalis*. (f-k) Correlation between flux and metabolite concentration (f-h) or between flux and enzyme concentration (i-k). Data within an organism (f,g,i,j) compares nutrient limited to batch conditions. Data across organisms (h,k) compares *S. cerevisiae* to *I. orientalis*. Each point represents median flux and concentration fold change for the pathway. Pearson’s R and p value are shown. Symbols in (h, k) are diamonds, CEN.PK in batch culture; circle, FY4 in batch culture; triangle, FY4 in nutrient limitation at 0.22 h^-1^. Black line shows slope = 1. Other pathways (folate, sugar, nucleic acid, lipid, amino acid) are plotted in different shades of grey.

The corresponding metabolomics and proteomics data provide a valuable resource for understanding the biochemical basis by which these fluxes are achieved, especially in the non-model yeast. For example, they can be assessed on a reaction-by-reaction basis to identify physiologically meaningful metabolic regulators ^44^ (Fig. S3a showing glyceraldehyde-3-phosphate dehydrogenase, or GAPD, as an example). We were able to identify allosteric regulation in 19 out of 51 examined reactions in *I. orientalis* (Supplementary Table). Seven of these regulations have also been reported in *S. cerevisiae*, including classical ones such as citrate inhibition of phosphofructokinase ^31^ and fructose-1,6-bisphosphate activation of pyruvate kinase ^45^. Our analysis also revealed multiple previously unappreciated regulatory interactions. For example, ATP inhibits *I. orientalis* GAPD, a novel interaction that we biochemically verified (Fig. S3, a-c). Overall, the integration of *in vivo* enzyme and metabolite concentrations via Michaelis-Menten kinetics explained the vast majority of flux variation across physiological conditions (Fig. 2d). This reflects enzyme concentration, active site occupancy, and allosteric regulation by metabolites collectively accounting for most yeast flux control, without the need to invoke other mechanisms like enzyme covalent modification or localization.

### Flux control by enzyme concentration

Cells contain extensive programs for regulating protein levels. Across nutrient conditions, however, we observed remarkably stable enzyme concentrations. In contrast, both metabolic fluxes and metabolite concentrations varied much more than enzymes (Fig. 2c, Fig. S2a). For some reactions, enzyme levels even show negative correlation with flux (negative Pearson’s R in Fig. 2d with enzyme only). We assessed the extent of physiological flux control residing in enzymes and metabolites based on their metabolic leverage, the product of their physiological concentration variation across conditions and their flux control coefficient based on the best-supported kinetic model from the above quantitative analysis of physiological metabolic regulation. Across 51 evaluable reactions, we found that, across nutrient conditions, metabolites exert much more metabolic leverage than enzymes (Fig. 2e, Fig. S3d).

Indeed, within both yeast species, flux changes across physiological conditions correlate better with pathway substrate concentration changes than pathway enzyme concentration changes (based on median of fold change across pathway components) (Fig. 2, f-i). The maintenance of enzyme concentrations with reduced growth and metabolic flux suggests substantial spare enzyme capacity, which may facilitate rapid growth acceleration when nutrient conditions improve ^26,27,46,47^.

In contrast to metabolite-dominant flux control within each yeast in response to changing nutrient environment, flux differences between *S. cerevisiae* and *I. orientalis* strongly aligned with enzyme concentrations (Fig. 2, j-k). Given the 200 million years of evolution separating these two species ^34^, we expected that there might be substantial differences in enzyme properties that change metabolic flux between the two organisms. Highly expressed proteins, including central metabolic enzymes, however remained strongly conserved between the two yeasts at the protein sequence level (Fig. S4). Correspondingly, enzyme abundances account for a large fraction of flux variation, including the greater glycolysis flux in *S. cerevisiae* and faster TCA turning and oxidative phosphorylation in *I. orientalis* (Fig. 2k). Thus, within the tested yeast species, flux is predominately regulated at the level of metabolite concentrations and active site occupancy. In contrast, flux differences between these yeast species is predominately explained by enzyme concentrations.

### Proteome-efficiency of ATP generation

We next examined overall proteome allocation of both yeasts with absolute proteomic quantification calibrated by UPS2 standard, and found that metabolic genes (enzymes, transporters, and mitochondrial proteins) together accounted for just over half of the proteome in both species, with the other major proteome sectors being translation (mostly ribosomes) and transcriptional machinery (Fig. 3a). The fractional proteome allocation to these three major sectors was nearly identical across the two yeasts. The major difference was within the metabolic sector, which had three major components: anabolic, glycolytic, and respiratory (the latter being composed of mitochondrial, TCA, and oxidative phosphorylation proteins). The anabolic enzymes constituted a similar proteome fraction in both species, but there was a major reallocation between the two other sectors: *S. cerevisiae* expresses more glycolytic proteins, and *I. orientalis* more respiratory proteins.

**Figure 3.**
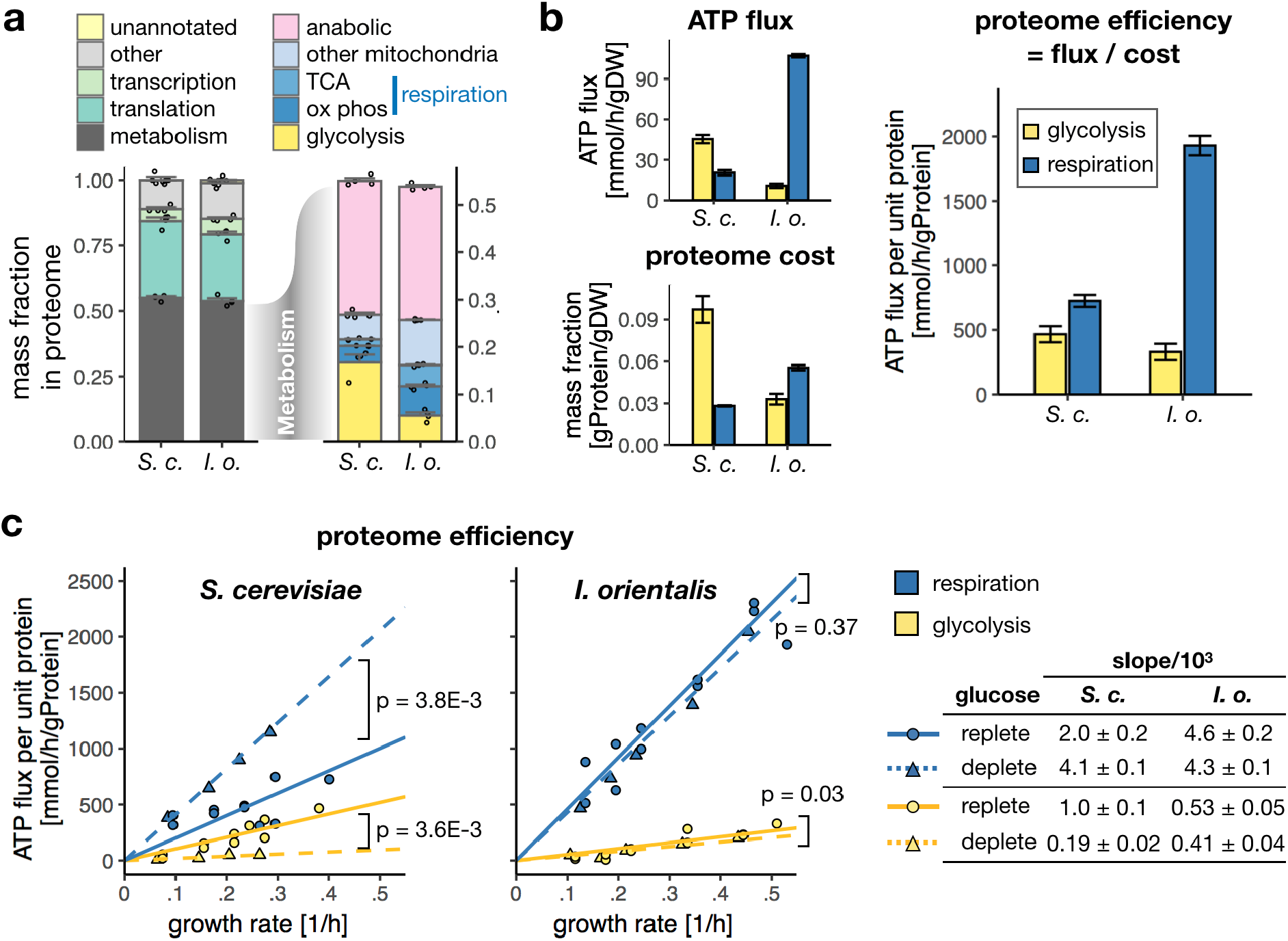
Proteome efficiency across nutrient conditions in *S.cerevisiae* and *I. orientalis*. (a) Proteome allocation of *S. cerevisiae* (CEN.PK) and *I. orientalis* in exponentially growing batch culture. Mean ± SE, n = 4 biological replicates. (b) ATP fluxes (from ^13^C-informed MFA), proteome mass fraction (of whole cell dry weight), and proteome efficiency for glycolysis and respiration in exponentially growing aerobic glucose-fed batch culture. ATP fluxes are shown as mean ± SE based on ^13^C-MFA confidence interval. Proteome efficiency is shown as mean ± SE, with error propagated from flux and proteome fraction measurements. (c) Proteome efficiency of respiration and glycolysis across different nutrient conditions in *S. cerevisiae* (left) and *I. orientalis* (right). For raw data, see Fig. S5. Solid line shows linear regression in glucose replete conditions (Batch, N-limit, and P-limit). Dashed line is glucose-depleted conditions (C-limit). P values are from ANOVA of linear model.

Together, our absolute proteome quantitation and flux analyses enabled quantification of ATP production per protein mass (i.e. proteome efficiency) of glycolysis and respiration in both species. We obtained the ATP flux from the genome-scale model, which included a mechanistic ratio of 3 ATP produced for every 10 protons translocated by ATP synthase ^48^. Including both TCA and oxidative phosphorylation proteins (but not other mitochondrial proteins) as the proteome cost of respiration, we found that, in batch culture, respiration is more proteome-efficient than glycolysis in both yeasts (Fig. 3b).

Despite comparing favorably to glycolysis, respiration in glucose-rich batch-cultured *S. cerevisiae* was much less proteome-efficient than in *I.orientalis*. Besides the absence of proton-pumping complex I in *S. cerevisiae*, we hypothesized that this also reflects spare respiratory capacity when glucose is abundant. Consistent with this, the proteome-efficiency of *S. cerevisiae* respiration increased (and of glycolysis fell) under glucose limitation (Fig. 3c, Fig. S5). In contrast, since *I. orientalis* defaults to respiration even when glucose is abundant, proteome-efficiency was unaffected by glucose availability.

Part of glycolytic and TCA flux is diverted to biosynthesis, and glycolytic flux is needed to generate respiratory substrate. We further assessed proteome efficiency of ATP generation of fermentation (turning glucose to ethanol) and respiration (turning glucose to CO2) by calculating a flux-partitioned proteome cost ^2^, which counts glycolytic proteins in the cost of respiration and discounts flux diverted to other pathways. In both yeasts, this flux-partitioned analysis identified respiration as the more proteome-efficient ATP production pathway (Fig. S6, a-b). Similarly, even if counting all mitochondrial proteins into respiration’s proteome cost, respiration remains more proteome-efficient than glycolysis in *I. orientalis* and glucose-limited *S. cerevisiae* (Fig. S6, c-d).

### Benefit of aerobic glycolysis

In *S. cerevisiae*, rate-yield tradeoff was believed to underlie the switching to carbon-inefficient fermentation at faster growth ^14–17^. Our data reveals, however, that respiration is both more energy- and proteome-efficient than glycolysis. Such efficiency would be expected to lead to greater fitness.

Consistently, *I. orientalis* outcompetes *S. cerevisiae* in co-culture under conditions requiring respiratory ATP production (ethanol, glucose limitation). Importantly, however, it also outcompetes under conditions where fermentation is a viable strategy (abundant glucose, nitrogen limitation, phosphorus limitation, and even sucrose as the sole carbon source, which *S. cerevisiae* can ferment on while *I. orientalis* alone cannot metabolize and presumably takes in glucose and fructose liberated by *S. cerevisiae*) (Fig. 4a, Fig. S7).

**Figure 4.**
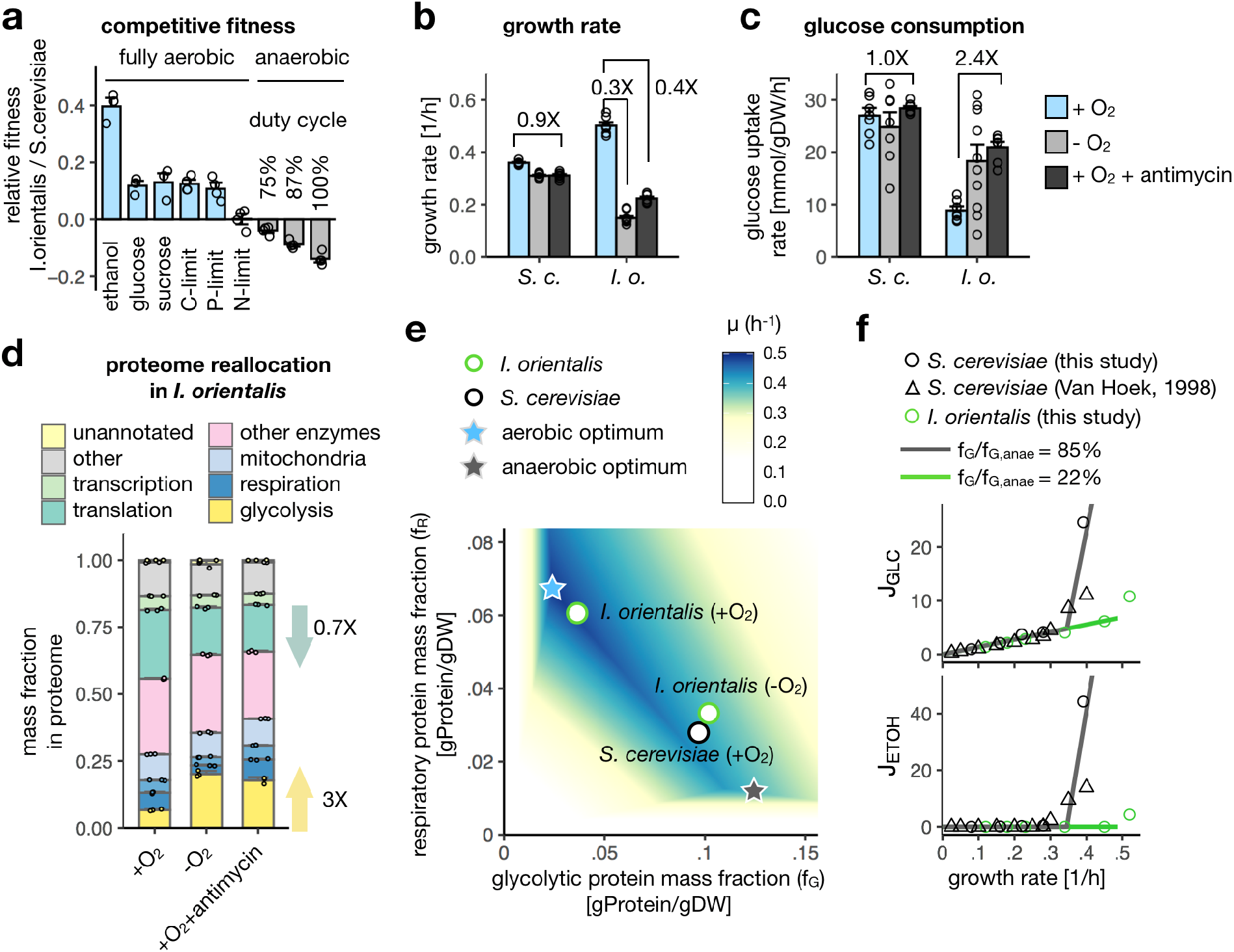
Aerobic glycolysis emerges from anaerobically primed proteome. (a) Fitness of *I. orientalis* and *S. cerevisiae* measured in competitive co-culture. Relative abundance of the two yeasts was measured by qPCR at 4 to 6 time points and used to obtain fitness. See method for details. Mean ± SE, n = 3 or 4 biological replicates. (b-c) Specific growth rates (b) and glucose consumption rates (c) for batch-cultured *S. cerevisiae* and *I. orientalis* with oxygen (blue), without oxygen (light grey), and with 10 μM antimycin (dark grey). Mean ± SE, n = 6 or 7 biological replicates. (d) Proteome allocation in *I. orientalis* in above conditions. Mean ± SE, n=3 biological replicates. Arrows show fold change in translational and glycolytic proteome compared to aerobic condition. (e) Respiro-fermentative growth rate (μ) was predicted from a proteome-constrained coarse-grained model parameterized with proteome efficiency measured from *S. cerevisiae*. Glycolytic (f_G_) and respiratory proteome fraction (f_R_), are mass fractions of whole cell dry weight. Optimal proteome fractions in aerobic and anaerobic condition were indicated as stars. Measured proteome fractions in glucose-fed batch cultures of *I. orientalis* (aerobic, +O_2_; or anaerobic, −O_2_) and *S. cerevisiae* (aerobic, +O_2_) are shown in circles. (f) Experimental glucose consumption (J_GLC_) and ethanol excretion (J_ETOH_) rates (symbols, in mmol/h/gDW) and prediction from proteome-constrained model (lines) under high (*S. cerevisiae*) or low (*I. orientalis*) glycolytic capacity (f_G_ relative to its anaerobic optimum, f_G,anae_). Literature data was obtained from Van Hoek 1998 ^32^.

If respiration is both more energy- and proteome-efficient than glycolysis, why does aerobic glycolysis occur? One possibility is the production of a toxic product that impairs competitors: ethanol ^49,50^. Another is carbon resource competition, essentially quick uptake of glucose ^1,33,51^. These might help *S. cerevisiae* compete with bacteria, but did not against *I. orientalis* (Fig. 4a).

We wondered whether the benefit of aerobic glycolysis might instead not be during aerobic growth, but rather in hedging for oxygen limitation (hypoxia) ^13^. *S. cerevisiae* was repeatedly exposed to oxygen limitation during human baking and winemaking ^52^. But oxygen limitation can also occur readily naturally, due to oxygen’s limited solubility (c_o2_ ≈ 230 μmol/L) and slow diffusion in water (D ≈ 2.3×10^-9^ m^2^/s). We measured the dissolved oxygen at the bottom of unstirred *S. cerevisiae* and *I. orientalis* cultures, and found that oxygen depletion occurred at relatively low culture density (Fig. S8). Notably, *S. cerevisiae* outcompeted *I. orientalis* in fully or cyclically oxygen-depleted co-cultures (Fig. 4a, Fig. S7). Consistently, we observed about 60% growth rate reduction in *I. orientalis* but not *S. cerevisiae* upon oxygen depletion or pharmacological inhibition of electron transport chain complex III by antimycin (Fig. 4b). Both oxygen depletion and antimycin increased glucose uptake rate by more than two-fold in *I. orientalis* (Fig. 4c), which is mediated by about 3-fold higher expression of glycolytic proteins (Fig. 4d). Notably, this glycolytic protein expression came at the expense of ribosomal proteins, consistent with the growth defect in anaerobic *I. orientalis* (Fig. 4d, Fig. S9).

### Proteomic hedging and aerobic glycolysis

Both *I. orientalis* and *S. cerevisiae* can tailor their respiratory versus glycolytic enzyme expression to environmental conditions. But this tailoring is incomplete: *I. orientalis* partially retains respiratory enzyme and mitochondrial protein expression in hypoxia (Fig. 4d), while *S. cerevisiae* retains high glycolytic enzymes in aerobic conditions (Fig. 2g).

To explore the consequences of incomplete proteome tailoring, we assembled a coarse-grained quantitative model of yeast growth and metabolism, where growth is limited both by ATP generation (fermentative or respiratory) and by translational machinery, jointly constrained by proteome capacity (Fig. 4e and Extended Data Note). Growth optimization is performed to find the optimal respiratory and glycolytic proteome allocation (f_R_ and f_G_, respectively). This minimal model captures the proteome tradeoff between optimal aerobic and anaerobic growth (Fig. 4e). While optimal aerobic growth is achieved via respiratory ATP production, when the proteome is constrained to always contain enough glycolytic enzyme for rapid anaerobic growth, aerobic glycolysis emerges as an optimal strategy (Fig. 4f).

### Mammalian ATP generation

We were curious if the greater proteome efficiency of respiration generalizes from yeast to mammals. To investigate this, we quantified the proteome efficiency of glycolysis and respiration in cultured cancer cells and mouse tissues (Fig. 5). ATP flux was from previous reports or estimated based on reported oxygen consumption rates ^54,55^ (Fig. 5a). The proteome fraction allocated to glycolysis and respiration was computed from published proteomics data ^56,57^, which shows that mouse tissues in general have greater respiratory proteome capacity like in *I. orientalis*, whereas cancer cell lines have more glycolytic proteome like in *S. cerevisiae* (Fig. 5b). Overall, the proteome efficiency of both glycolysis and respiration was lower in mammals than in yeast, consistent with mammals being under less stringent selection for proteome efficiency. Nevertheless, in both cultured cancer cells and *in vivo* tissues, respiration was the more proteome-efficient ATP generation pathway (Fig. 5c). Thus, in both yeast and mammals, mitochondrial respiration is more proteome-efficient than glycolysis.

**Figure 5.**
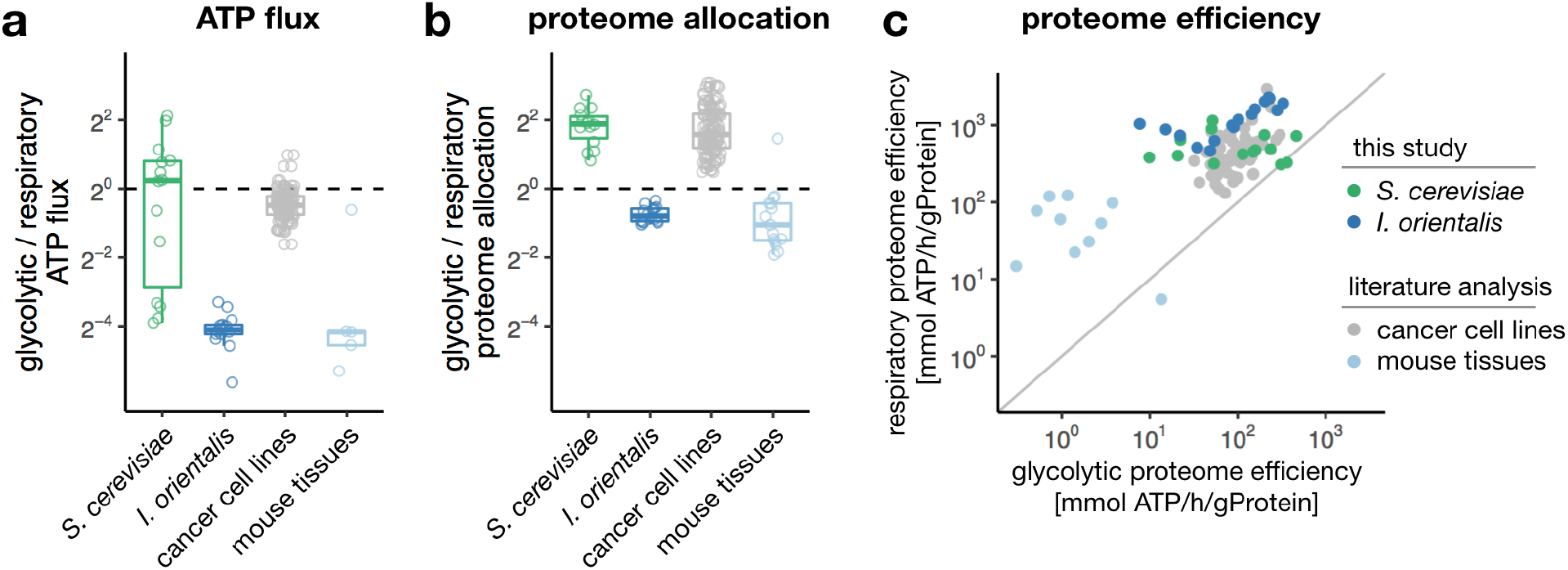
Respiratory ATP production is more proteome efficient than glycolysis in mammals. (a) Ratio between glycolytic and respiratory ATP production in *I. orientalis, S. cerevisiae*, NCI60 cancer cell lines, and mouse tissues. For yeasts, each data point represents a nutrient condition. For cancer cell lines and mouse tissues, each point represents an individual cell line or tissue. (b) As in (a), for proteome allocation. (c) Corresponding glycolytic versus respiratory proteome efficiency.

## Discussion

Here we report in-depth proteomic, metabolomic, and metabolic flux characterization of two divergent budding yeasts across various environmental conditions. The resulting data reveal principles of yeast metabolism and its regulation. Prior integrative ‘omic analysis of *E. coli* and *S. cerevisiae* concluded that metabolism is substantially ‘self-regulated’, i.e. that changes in metabolic flux are caused more by metabolites themselves than transcriptional and translational reprogramming of enzyme levels ^44,58–60^. This conclusion is reinforced by analogous analysis of *I. orientalis* here, which shows even less proteome variation across most environmental conditions than *S. cerevisiae*, and yet greater dominance of metabolic flux control by metabolites themselves.

In contrast to the limited impact of the proteome on flux control within each species, across the two species, metabolic differences are mainly encoded by protein abundances. Given that these two yeasts diverged roughly 200 million years ago, the ability to explain most of their metabolic differences through enzyme concentrations – rather than changes in the properties of enzymes themselves – is notable, and speaks to the importance of proteome allocation in driving metabolic divergence even across long timescales ^61^.

The most striking metabolic difference between *I. orientalis* and *S. cerevisiae* is that, in the presence of abundant glucose, the former respires while the latter engages in aerobic glycolysis. We show that, across a wide range of aerobic conditions, the more respiratory yeast grows faster and has superior competitive fitness. This aligns with respiration requiring less of the cell’s precious proteome capacity to achieve the same growth-required ATP flux. Quantitative analysis of mammalian cancer cells and tissues demonstrates that respiration is also more proteome-efficient than glycolysis in mammals.

Prior careful evaluation of proteome efficiency in *E. coli* reached a seemingly opposite conclusion, finding that acetate fermentation is favored for its proteome efficiency ^2^. Acetate overflow metabolism in *E. coli*, however, involves a blend of glycolytic and respiratory ATP generation with 4 NADH feeding into the electron transport chain for each glucose. This provides an ATP yield of about 12 per glucose, the majority of which is made via the oxidative phosphorylation (compared to 2 ATP per glucose in yeast or mammalian aerobic glycolysis). Thus, aerobic ‘fermentation’ in *E. coli* is proteome efficient only because it generates substantial respiratory ATP.

Overall, supported by our data in *S. cerevisiae* and *I. orientalis*, we propose that cells of a given type tend to have a characteristic metabolic proteome that varies only modestly across conditions. In this nearly fixed enzyme network, changing substrate levels induce different fluxes, providing metabolic flexibility without the need for extensive proteome remodeling. A benefit of such proteome constancy is that cells are prepared in advance for changing metabolic environments. One of the most important metabolic fluctuations cells face is shifting oxygen availability ^62^. We thus posit that aerobic glycolysis occurs not because it is beneficial per se, but as a side effect of maintaining a fermentative proteome that effectively supports both aerobic and anaerobic growth.

## Acknowledgement

We thank the members of Rabinowitz lab for discussion on experiments and the manuscript; S. Silverman and J. Avalos for yeast strains, L. Ryazanova for help with proteomics experiment, P. F. Suthers for discussion on the genome-scale model, M. Gupta for discussion on protein regulation, N. Piyush and Z. Zhang for advice on competitive fitness, and Z.-Y. Wu for experimental support. This work was funded by DOE grant DE-SC0018260 and the DOE Center for Advanced Bioenergy and Bioproducts Innovation (Award Number DE-SC0018420). Any opinions, findings, conclusions or recommendations expressed in this publication are those of the author(s) and do not necessarily reflect the views of the U.S. Department of Energy.

## Author contributions

Y. S., M.W. and J.D.R. designed this study. Y.S. performed most of the experiments and data analysis. H.V.D. designed and performed genome-scale metabolic flux analysis with the input from Y.S., J.I.H., and C.D.M. E.C., H.B., and A.S. performed proteomics measurement. C.M.C. performed nutrient limited culture and measurement. R.P.R. designed and performed enzyme purification and qPCR. J.P. performed enzyme purification and competitive growth experiments.

Z. F. created mutant yeast strains with the input from H.Z., and contributed to enzyme purification.

S. D. and Y.Y. contributed to yeast growth measurement. V.T. contributed to enzyme purification.

T. X. contributed to metabolomics measurements. D.W. contributed to enzyme constrained modeling. L.Y. contributed to oxygen consumption measurement. Y.S. and J.D.R. wrote the manuscript with input from all co-authors.

