## supplementary information for "Proteome capacity constraints favor respiratory ATP generation"

### Extended Data and Note for
### Proteome capacity constraints favor respiratory ATP generation

Yihui Shen [1,2], Hoang V. Dinh [5], Edward Cruz [2,4], Catherine M. Call [1,2], Heide Baron [2], Rolf-Peter Ryseck [1,2], Jimmy Pratas [1,2], Arjuna Subramanian [2], Zia Fatma [6], Daniel Weilandt [1,2], Sudharsan Dwaraknath [8], Tianxia Xiao [1,2], John I. Hendry [5], Vinh Tran [6], Lifeng Yang [1,2], Yasuo Yoshikuni [8], Huimin Zhao [6,7], Costas D. Maranas [5], Martin Wühr [2,4], Joshua D. Rabinowitz [1,2,3]

1 Department of Chemistry, Princeton University, Princeton, NJ, USA
2 Lewis Sigler Institute for Integrative Genomics, Princeton University, Princeton, NJ, USA
3 Ludwig Institute for Cancer Research, Princeton Branch, Princeton, NJ, USA
4 Department of Molecular Biology, Princeton University, Princeton, New Jersey 08544, USA
5 Department of Chemical Engineering, The Pennsylvania State University, University Park, PA, USA
6 Carl R. Woese Institute for Genomic Biology, University of Illinois Urbana-Champaign, Urbana, IL, 61801, USA
7 Department of Chemical and Biomolecular Engineering, University of Illinois at Urbana-Champagne, Urbana, IL, USA
8 US Department of Energy Joint Genome Institute, Lawrence Berkeley National Laboratory, Berkeley, CA 94720, USA

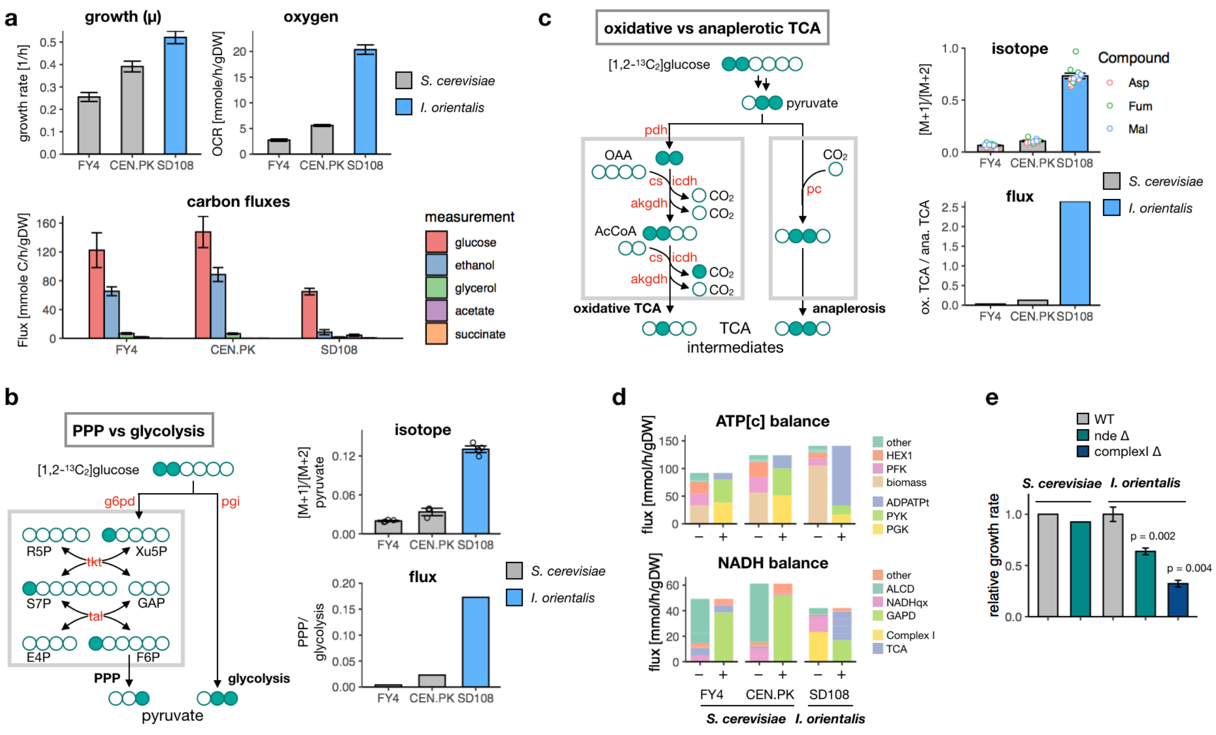

**Extended Data Fig. 1 Flux analysis in batch-grown S. cerevisiae and I. orientalis**

(a) Growth rate, oxygen consumption rate, and carbon metabolite fluxes were measured for *S. cerevisiae* (FY4 or CEN.PK strains) and *I. orientalis* SD108 grown in YNB with 20g/L glucose. Mean ± SE, n = 3 biological replicates.

(b) Isotopic signature in pyruvate reveals higher flux through pentose phosphate pathway (PPP) in *I. orientalis*. Left, carbon atom mapping of [1,2-$^{13}C_2$] glucose going through PPP or glycolysis, where filled circles represent $^{13}C$. Right top, measured [M+1]/[M+2] ratio of pyruvate from [1,2-$^{13}C_2$] glucose tracing (1:1 mixed with unlabeled glucose), mean and standard error of n = 4 biological replicates. Right bottom, flux split ratio between PPP and glycolysis calculated as (G6PD - PRPPS)/PGI flux determined from $^{13}C$ genome-scale MFA. G6PD, glucose-6-phosphate dehydrogenase; PRPPS, phosphoribosylpyrophosphate synthetase; PGI, phosphoglucose isomerase.

(c) Isotopic signature in TCA intermediates reveals significantly higher oxidative TCA activity in *I. orientalis*. Average [M+1]/[M+2] ratio from [1,2-$^{13}C_2$] glucose tracing (1:1 mixed with unlabeled glucose) is shown for three TCA metabolites (Asp, aspartate; Fum, fumarate; Mal, malate), mean and standard error of n = 3 or 4 biological replicates. Flux ratio between oxidative and anaplerotic TCA is calculated as (CS+AKGDH+SUCD)/3/PC. CS, citrate synthase; AKGDH, alpha-ketoglutarate dehydrogenase; SUCD, succinate dehydrogenase; PC, pyruvate decarboxylase.

(d) Consumption (-) and production (+) flux contributing to the balance of cytosolic ATP (ATP[c]) and whole-cell NADH. PFK HEX1 PYK PGK GAPD, reactions in glycolysis; ADPATPt, mitochondrial transporter for ADP and ATP; PYRDC, pyruvate decarboxylase, PYRtps; mitochondrial pyruvate transporter; NADHqx, external NADH quinone oxidoreductase; NADHcplx, electron transport chain complex I; ALCD, alcohol dehydrogenase; PDH AKGDH MDH, reactions associated with TCA. Fluxes are best estimate from genome-scale $^{13}C$ MFA.

(e) Growth impact of NADH dehydrogenase deletion in *S. cerevisiae* and *I. orientalis*. Data for *S. cerevisiae* is from a previous study[1]; data for *I. orientalis* is determined in this study and shows best estimate and standard error from exponential fitting.

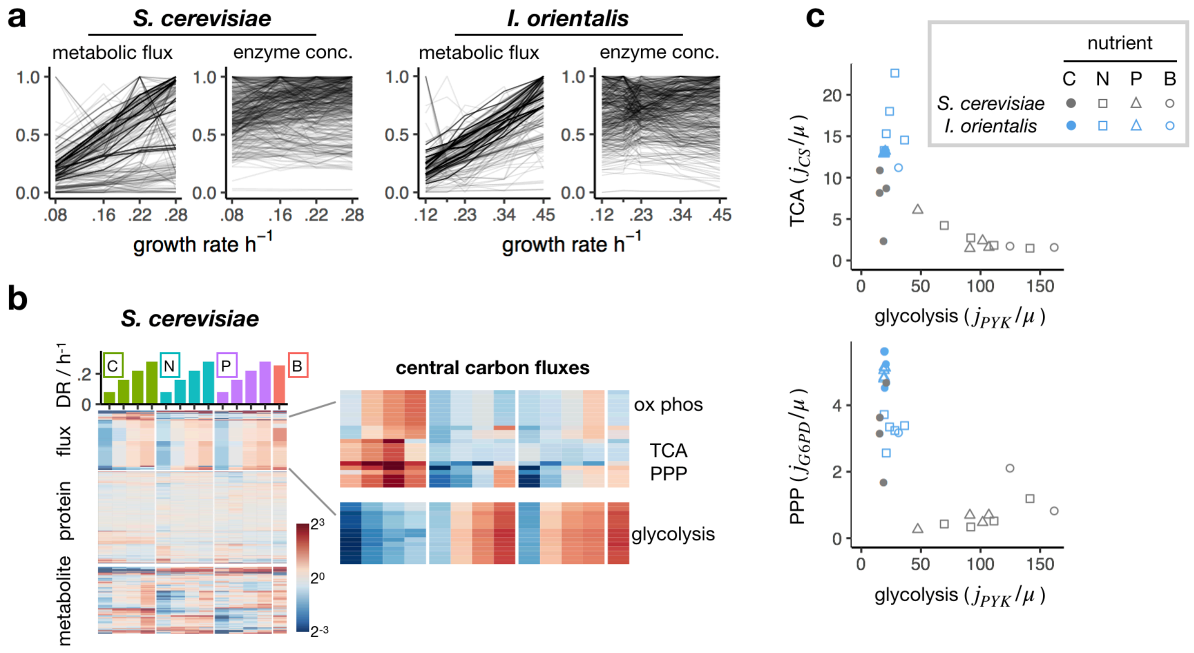

**Extended Data Fig. 2 Multi-omics data of *S. cerevisiae* and *I. orientalis* across different growth conditions**

(a) Dependence of metabolic flux and enzyme concentration on growth rate. Flux and enzyme concentration are normalized to maximum across nutrient conditions. For each reaction, a linear regression is done for flux ~ growth rate. % variance explained is calculated, and averaged across all reactions. 53% of flux variation in *S. cerevisiae* and 71% in *I. orientalis* can be explained by growth rate alone. 21% of enzyme variation in *S. cerevisiae* and 23% in *I. orientalis* can be explained by growth rate alone.

(b) Change in metabolic flux, protein abundance and metabolite concentration in *S. cerevisiae* across growth conditions. Each row represents a reaction, or a protein, or a metabolite, normalized to the geometric mean across all the conditions. 4 different dilution rates (DR) were used for each limiting nutrient. Limiting nutrient: C, carbon; N, nitrogen; P, phosphorus; B (batch), none.

(c) Metabolic phenotype of C-limited *S. cerevisiae* resembles *I. orientalis*. Fluxes through TCA, glycolysis, PPP are represented by flux through citrate synthase (CS), pyruvate kinase (PYK), and glucose-6-phosphate dehydrogenase (G6PD), respectively. Values are shown after normalization by growth rate ($\mu$).

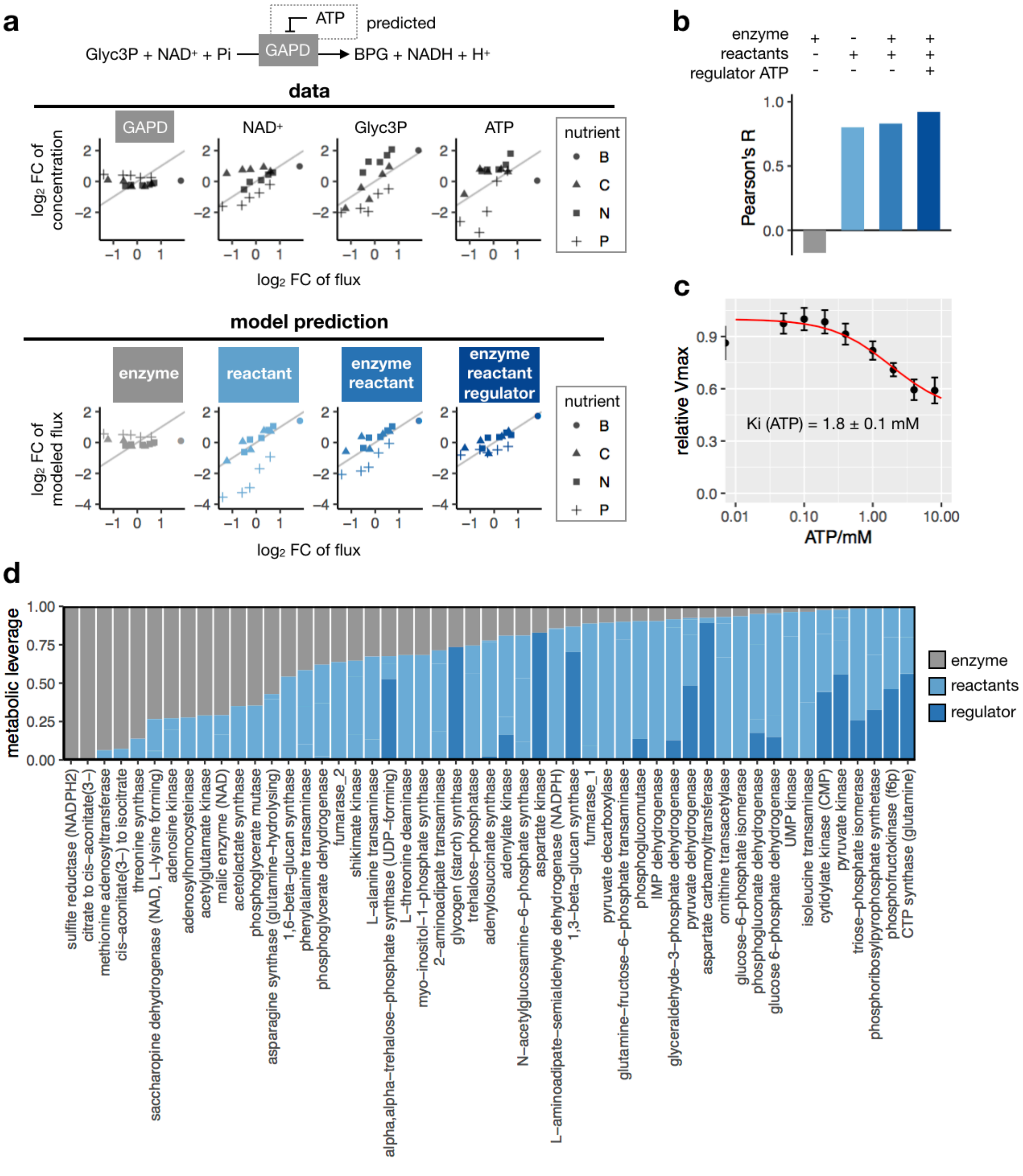

**Extended Data Fig. 3 Identification of physiologically relevant metabolic regulators in *I. orientalis***

(a) Multi-omics data integration, showing glyceraldehyde-3-phosphate dehydrogenase (GAPD) as an example. Top row shows relative fold change (FC) in enzyme abundance or metabolite concentration with respect to fold change in flux across different growth conditions in log2 scale. Limiting nutrient: C, carbon; N, nitrogen; P, phosphorus; B, none. Bottom row shows modeled flux accounting for enzyme alone, reactant alone, enzyme and reactants with or without regulator.

(b) Correlation between modeled flux change and experimental flux change in (a) quantified by Pearson's correlation coefficient.

(c) Biochemical validation of ATP inhibition of *I. orientalis* GAPD. The enzyme activity of recombinantly purified GAPD was measured with different ATP concentrations. Data show mean and standard error of n = 9 replicates, and are fitted to partial inhibition model $Vmax \sim 1/([ATP]+Ki) + 1-1/Ki$ to obtain $Ki = 1.8 \pm 0.1$ mM.

(d) Partition of metabolic control (same as Fig. 2e, shown here in larger format with reaction names).

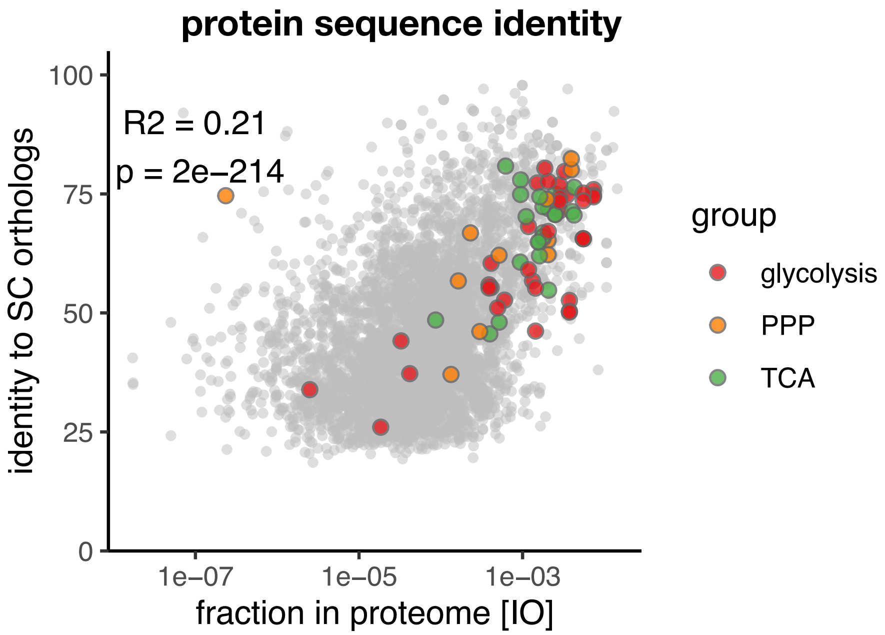

**Extended Data Fig. 4 Protein sequence identity between *S. cerevisiae* and *I. orientalis***

Genome-wide protein sequence identity is obtained through blastp between *S. cerevisiae* and *I. orientalis* orthologs, and shown with respect to protein abundance in *I. orientalis* (in log scale). Pearson's R = 0.21, p = 2E-214. Enzymes in glycolysis (red), PPP (orange), and TCA (green) are highlighted.

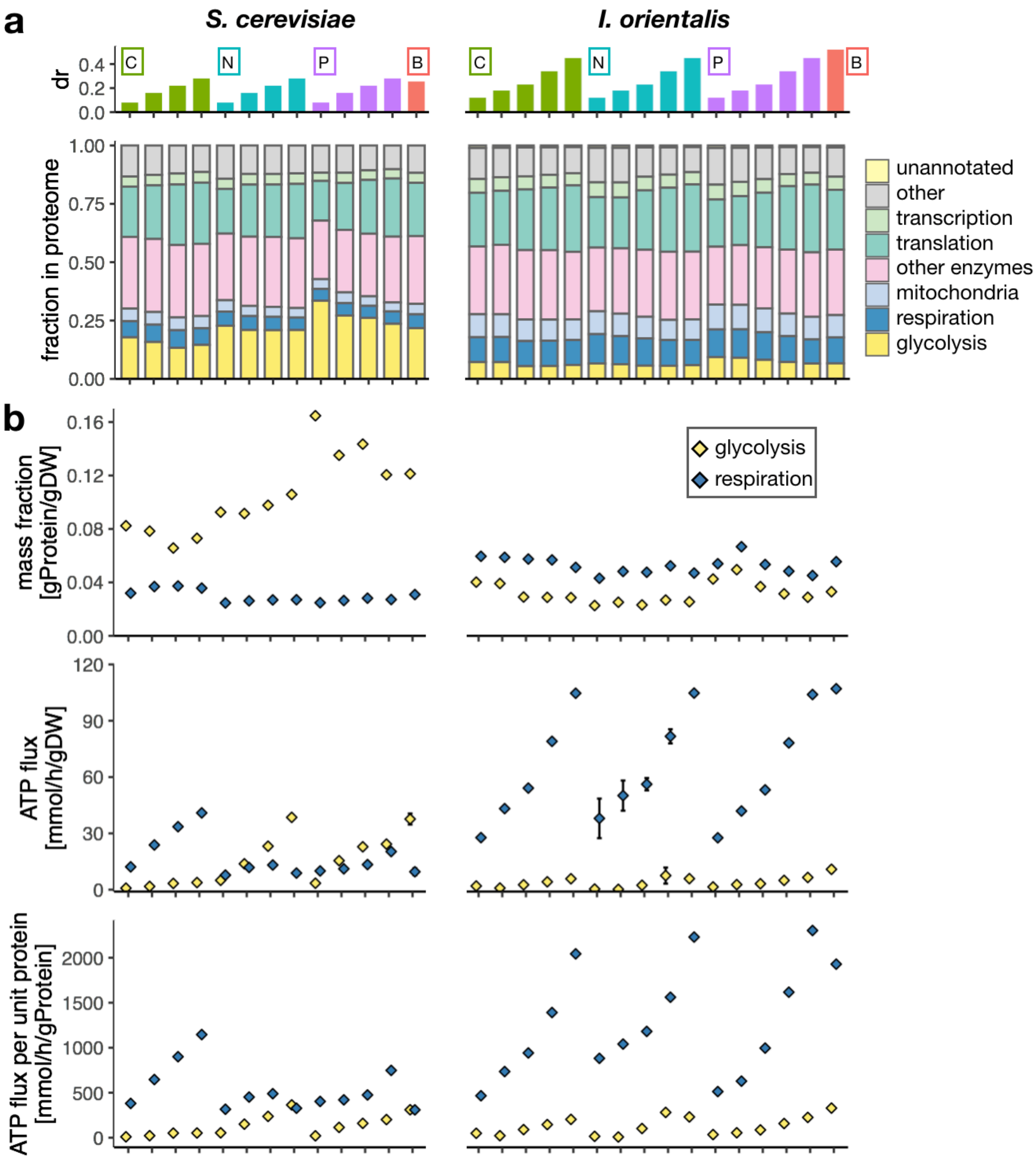

**Extended Data Fig. 5 Proteome allocation and ATP flux *S. cerevisiae* and *I. orientalis* across nutrient conditions**

(a) Proteome fraction of 7 functional sectors are shown for *S. cerevisiae* (FY4) and *I. orientalis* across nutrient conditions described in Fig. 2 and Fig. S2. n=1 for each condition.

(b) Proteome fraction (top), MFA-derived ATP fluxes (middle), and proteome efficiency of ATP production (bottom) for glycolysis and respiration are shown for conditions in (a), similar to Fig. 3b.

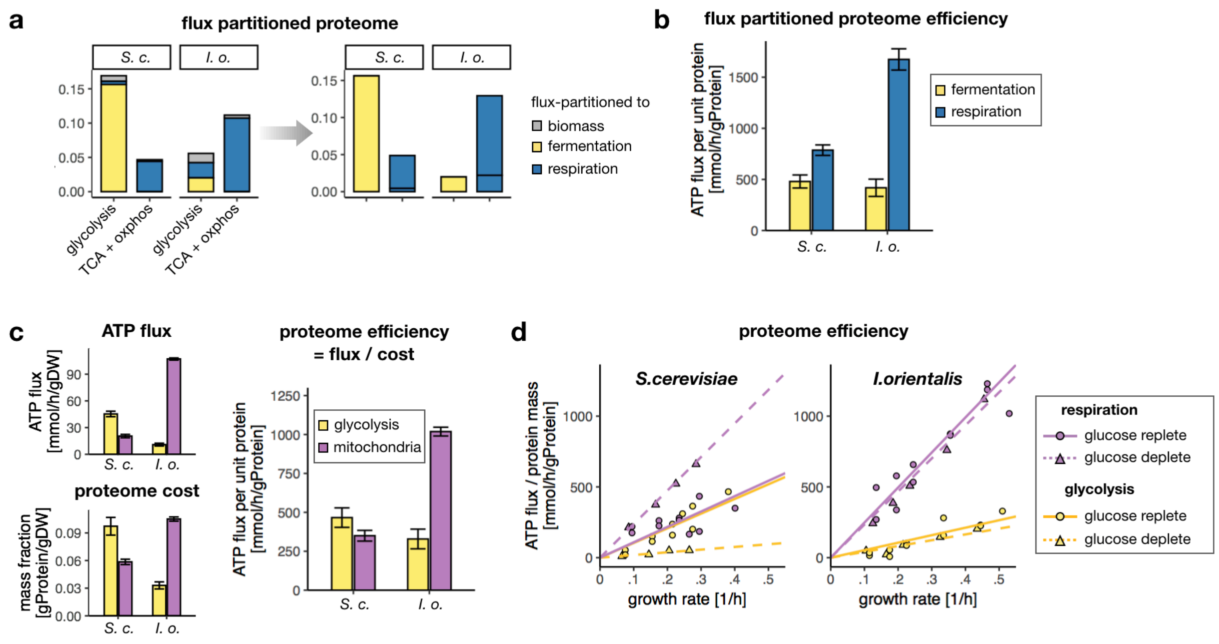

**Extended Data Fig. 6 Proteome efficiency accounting for flux partitioned proteome cost or non-enzymatic mitochondrial cost**

(a) Proteome allocation to respiration and fermentation (right) based on flux partitioning of the glycolytic and TCA and oxphos proteome (left).

(b) Proteome efficiency reevaluated from partitioned proteome cost in (a).

(c-d) Proteome efficiency in batch culture (c) or across all conditions (d), similar to Fig. 3 (b and c), except that mitochondrial proteome cost includes all mitochondrial proteins.

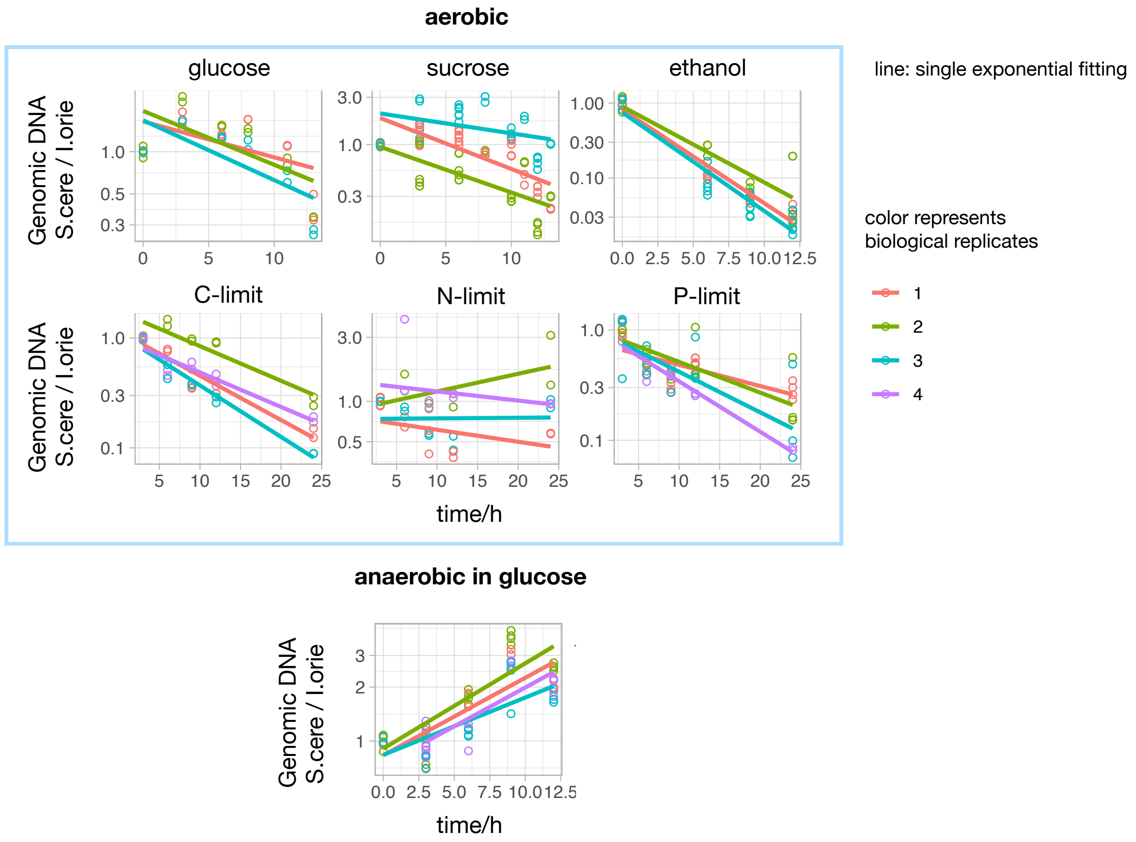

**Extended Data Fig. 7 Relative abundance of *S. cerevisiae* and *I. orientalis* in competitive co-culture**

*S. cerevisiae* (CEN.PK) and *I. orientalis* were inoculated at about same density in coculture with indicated nutrient conditions. Top row, yeast nitrogen base with 20g/L glucose or 20g/L sucrose or 10g/L ethanol, cultured aerobically. Middle row, continuous aerobic culture at $0.1h^{-1}$ dilution rate, with same nutrient limitation as described in Fig. 2. Bottom row, anaerobic culture with 20g/L glucose. Genomic DNA was extracted from the mixture and analyzed by qPCR to determine relative abundance between *S. cerevisiae* and *I. orientalis*. Color represents biological replicates (n = 3 or 4), each with n = 2 technical replicates. Solid line shows single exponential fitting.

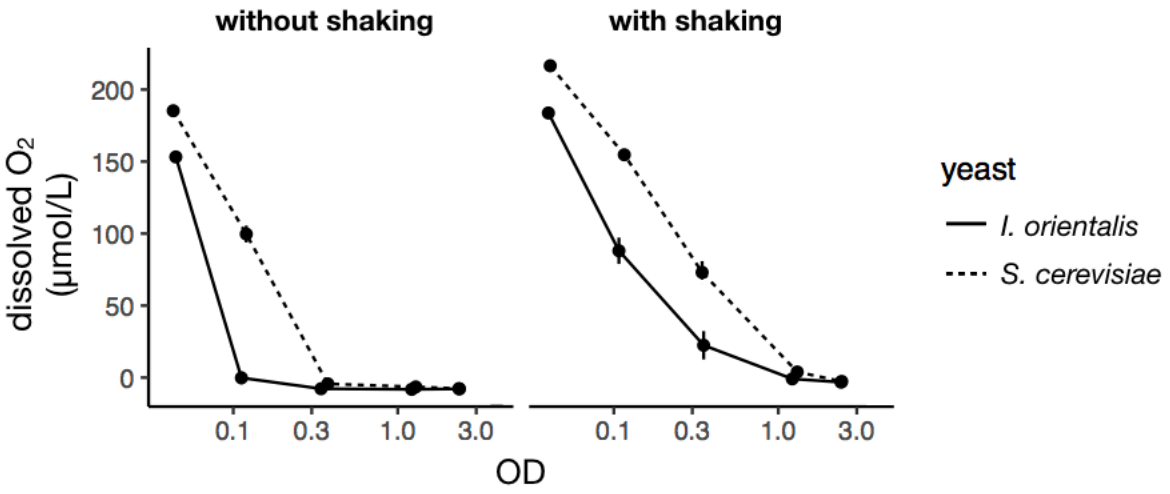

**Extended Data Fig. 8 Yeast deplete oxygen in liquid culture**

Dissolved oxygen measured at the bottom of the culture. Fresh culture was added to the plate at indicated density and allowed to adapt for 15min. Oxygen concentration was then measured with or without active shaking. The typical maximal OD from fully aerated culture using same media is about 4.

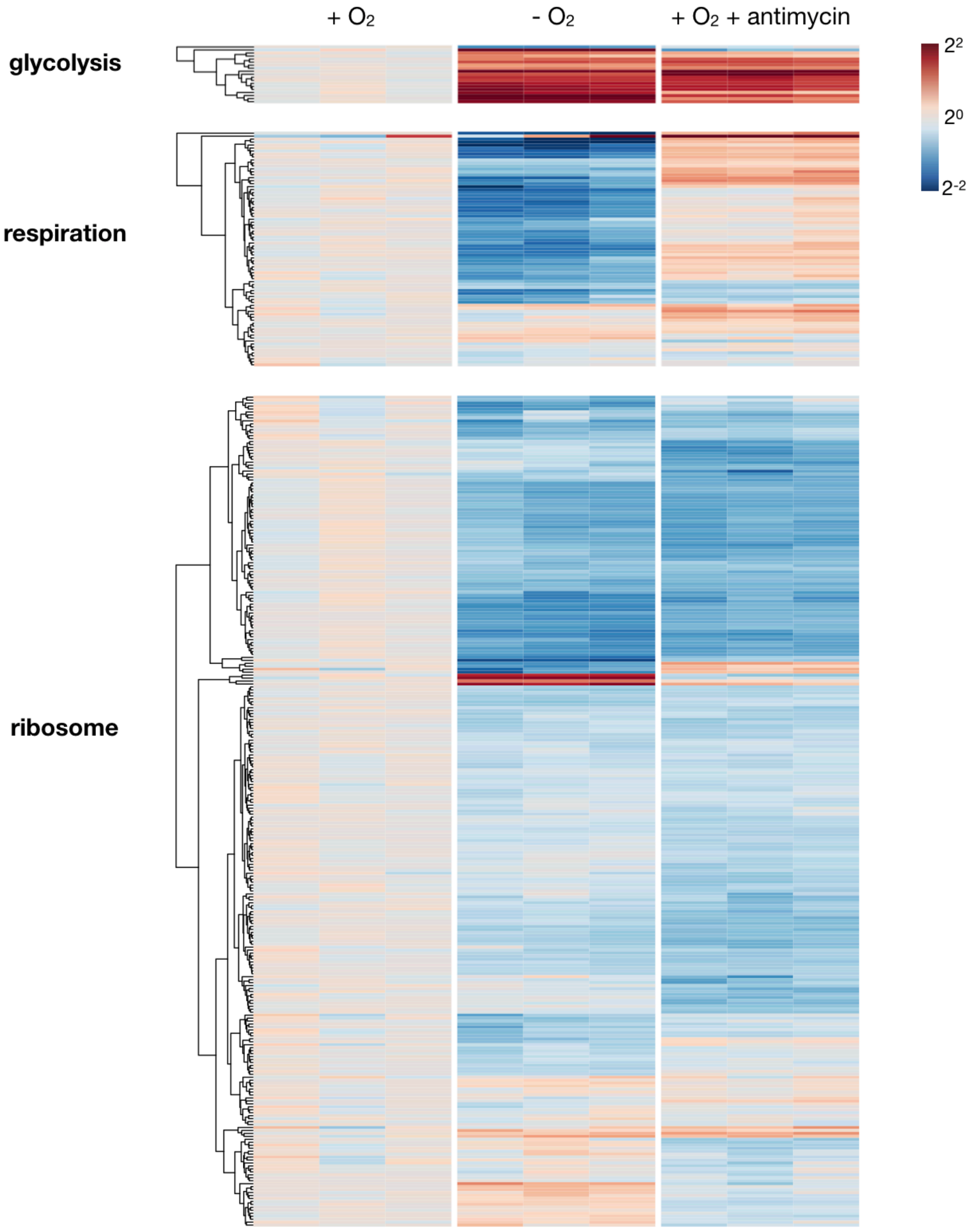

**Extended Data Fig. 9 Proteome remodeling in *I. orientalis* under respiratory deficiency**

Protein abundance in I. orientalis under aerobic (+$O_2$) or anaerobic (-$O_2$) conditions or aerobic with 10µM antimycin for 8hrs (+$O_2$+antimycin) on YNB and glucose. Fold change relative to the average of aerobic condition. n=3 biological replicates. Each row represents a protein grouped to glycolysis, respiration, and translation, and hierarchical clustered within each functional group.

**Extended Data Note - Coarse-grained proteome-constrained model**

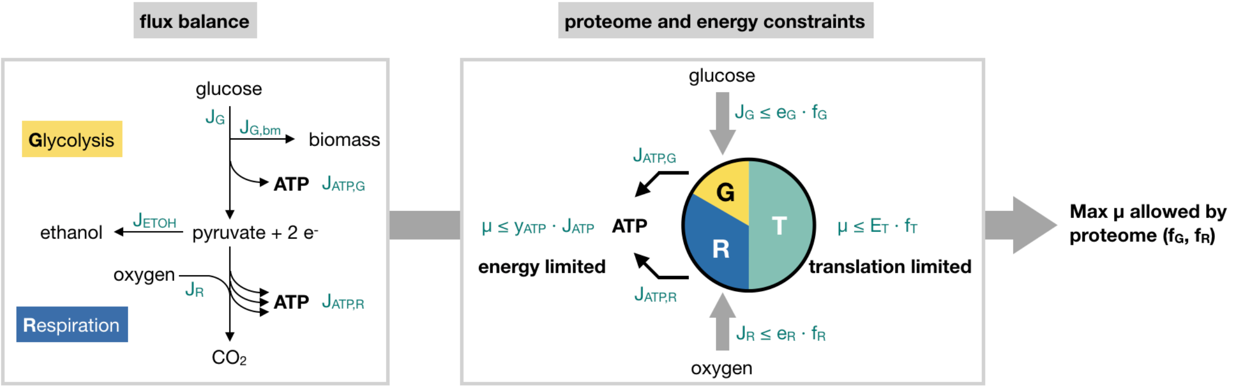

**Figure N1|** A coarse-grained model where yeast growth is constrained by flux balance, ATP availability, and translational capacity. G, glycolysis; R, respiration; T, translation; $\mu$, growth rate; J, flux; f, proteome fraction.

<u>Variables</u>

$\mu$, specific growth rate, 1/h

$f_X$, proteome mass fraction in whole cell dry weight for pathway $X$, in [gProtein/gDW]

$J_G$, glucose consumption rate, in [mmol glucose/gDW/h]

$J_R$, respiration rate, or oxygen consumption rate, in [mmol O/gDW/h]

$J_{ETOH}$, ethanol excretion rate, in [mmol EtOH/gDW/h]

<u>Parameters</u>

PO, P/O ratio, i.e. ATP synthesized per O atom, unitless. See **'obtain parameters from genome-scale model or experimental data'**.

$y_{ATP}$, growth yield per ATP, in [gDW/mmol ATP].

$e_X$, proteome efficiency of pathway X (X = G, R), in unit of [mmol/h/gProtein], defined as flux [mmol/h/gDW] per protein mass fraction of whole cell dry weight [gProtein/gDW]. It is the pathway equivalent of specific enzyme activity.

$E_T$, translational proteome efficiency, in unit of [gDW/h/gProtein], defined as growth rate [1/h] per translational protein mass fraction of whole cell dry weight [gProtein/gDW]. It is equivalent to the product of protein translation rate, $e_T$ [mmol AA/h/gProtein], and protein synthesis required for biomass renewal $y_{protein}$ [gDW/mmol AA]. ($J_{protein} \le e_T * f_T$, $\mu = y_{protein} * J_{protein}$. Define $E_T = y_{protein} * e_T$, thus $\mu \le E_T * f_T$.)

$f_0$, total proteome fraction allocated to energy and translation, in [gProtein/gDW].

**Carbon flux balance**

Flux balance for glucose (Fig. N1), $J_G = J_{G,bm} + J_{G,fermentation} + J_{G,respiration}$. $J_{G,bm}$, glucose equivalent ending up in biomass, in [mmol glucose/gDW/h], is proportional to growth rate, with a linear coefficient s [mmol glucose/gDW], i.e.

$$J_{G,bm} = \mathrm{s} * \mu \quad \text{[E1]}$$

According to reaction stoichiometry: fermentation, Glucose → 2 EtOH + 2 $CO_2$ (assuming ethanol to be the major waste product) and respiration, Glucose + 12 [O] → 6 $CO_2$ + 6$H_2O$, we have $J_{G,fermentation} = \frac{1}{2} \cdot J_{ETOH}$, and $J_{G,respiration} = \frac{1}{12} \cdot J_R$, and thus

$$J_G = J_{G,bm} + \frac{1}{2} \cdot J_{ETOH} + \frac{1}{12} \cdot J_R \quad \text{[E2]}$$

Ethanol overflow flux, from [E1] [E2]

$$J_{ETOH} = 2 * (J_G - s * \mu - J_R/12) \quad \text{[E3]}$$

Each glucose through glycolysis generates 2 ATP molecules, and since $J_{G,bm}$ does not lead to energy production, ATP flux from glycolysis is (with [E1])

$$J_{ATP,G} = 2 * \left(J_G - J_{G,bm}\right) = 2 * (J_G - s * \mu) \quad \text{[E4]}$$

ATP flux from respiration is $J_{ATP,R} = \mathrm{PO} * J_R$. Therefore, with [E1], total ATP flux can be derived from $J_G$ $J_R$ and $\mu$.

$$J_{ATP} = J_{ATP,G} + J_{ATP,R} = 2 * (J_G - s * \mu) + \mathrm{PO} * J_R \quad \text{[E5]}$$

**Optimal growth rate under energy and proteome constraint**

For each proteome configuration $(f_G, f_R)$, find the maximal allowed $\mu$ subject to the following constraints.

Proteome capacity constraints

Glycolytic capacity $J_G \leq e_G * f_G$ [E6]

Respiratory capacity $J_R \leq e_R * f_R$ [E7]

Translational capacity $\mu \leq E_T * f_T$ [E8]

Proteome constraint $f_G + f_R + f_T = f_0$ [E9]

From [E5] [E6] [E7] we could obtain **energy-limited growth**

$$\mu \leq y_{ATP} * J_{ATP} = y_{ATP} * (2 * (J_G - s * \mu) + \mathrm{PO} * J_R) \quad \text{[E10]}$$

From [E8] [E9] we could obtain **translation-limited growth**

$$\mu \leq E_T * (f_0 - f_G - f_R) \quad \text{[E11]}$$

In addition to [E6] [E8], $J_G$ and $J_R$ can also be constrained according to nutrient condition, eg. $J_G$ can be constrained to measured glucose uptake rate in carbon limited continuous culture, whereas $J_R = 0$ in anaerobic culture. A non-negative constraint $J_{ETOH} \geq 0$ is also applied to all conditions.

A minimal respiratory proteome is required to generate anabolic precursors like α-ketoglutarate, hence

$$f_R \geq \beta * \mu \tag{E12}$$

where $\beta$ [gProtein/gDW·h] is the minimal respiratory proteome mass fraction [gProtein/gDW] per unit growth rate [1/h].

**Obtain parameters from genome-scale model or experimental data**

(1) PO = 1.8, using theoretical value from ATP synthase which produces 3 ATP molecules for every 10 protons translocated, same in the *S. cerevisiae* genome-scale model. Relevant reactions are shown below with their id's in genome-scale model if available ($H_2O$ is omitted for simplicity).

| | |
|---|---|
| glucose → 5 NADH + 1 FADH + 6 $CO_2$ | complete oxidation of glucose |
| (N,F)ADH + [O] + 6 H_m → (N,F)AD + 6 H_c | electron transport chain |
| 3 H_m + 3 ADP_m + 3 Pi_m + 10 H_c → 3 ATP_m + 10 H_m | ATPS_m |
| 3 H_c + 3 Pi_c → 3 H_m + 3 Pi_m | PItps_m |
| 3 ATP_m + 3 ADP_c → 3 ATP_c + 3 ADP_m | ADPATPt_c_m |
| 3 ATP_c → 3 ADP_c + 3 Pi_c + 3 H_c | ATP hydrolysis |

(2) $s = 8.22$ [mmol glucose/gDW], obtained from fitting experimental $(J_G - J_{ATP,G}/2) \sim \mu$ according to [E4]. In carbon-limited conditions, this is about half of total glucose consumption, $J_G/\mu \approx 14.1$ [mmol glucose/gDW].

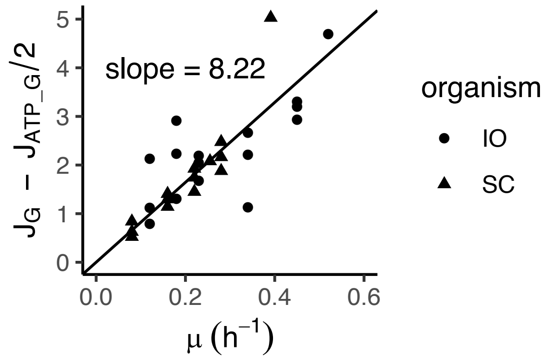

**Figure N2**| Determining glucose consumption directed to biomass ($s$) by linear fit between ($J_G - J_{ATP,G}/2$) and $\mu$.

(3) $y_{ATP}$ = 6.13×10$^{-3}$ [gDW/mmol ATP], obtained from fitting experimental $\mu \sim J_{ATP}$ (based on [E5] and experimental $J_G$ and $J_R$) from *S. cerevisiae*. $y_{ATP}^{-1}$ = 163 [mmol ATP/gDW], the majority of which is contributed by the growth-associated ATP maintenance (GAM). For reference, in the genome-scale model of *S. cerevisiae*, GAM = 139 [mmol ATP/gDW]. The difference from $y_{ATP}^{-1}$ is that GAM does not include ATP consumed for metabolite biosynthesis.

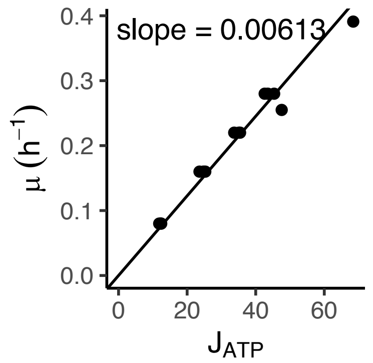

**Figure N3**| Determining growth yield per ATP, $y_{ATP}$, by linear fitting $\mu$ [h$^{-1}$] to $J_{ATP}$ [mmol/h/gDW].

(4) $e_G$ = 285 [mmol/h/gProtein], obtained from finding the max$(J_G/f_G)$ among all conditions for *S. cerevisiae*, based on [E6]. 2*$e_G$ is approximately the glycolytic proteome efficiency for ATP production defined in Fig. 3b.

(5) $e_R$ = 608 [mmol/h/gProtein], obtained from finding the max$(J_R/f_R)$ among all conditions for *S. cerevisiae*, based on [E7]. PO*$e_R$ is approximately the respiratory proteome efficiency for ATP production defined in Fig. 3b.

(6) $E_T$ = 2.38 [gDW/h/gProtein], obtained from finding the max$(\mu/f_R)$ among all conditions for *S. cerevisiae*, based on [E8]

(7) $f_0 = 0.30 \pm 0.02$ [gProtein/gDW], obtained from absolute proteomics in batch-cultured *S. cerevisiae*.

(8) $\beta = 0.030$ [gProtein/gDW·h], obtained from min $\left(\frac{\frac{J_{PDH\_m} - J_{AKGDH\_m}}{J_{PDH\_m}} * f_R}{\mu}\right)$ among all conditions for *S. cerevisiae* based on (Eq.7), where $\frac{J_{PDH\_m} - J_{AKGDH\_m}}{J_{PDH\_m}}$ represents the fraction of respiratory proteome used for anabolism. This minimum occurs in fully respiratory growth with carbon limitation, where anabolism accounts for less than a quarter of respiratory proteome usage ($\frac{J_{PDH\_m} - J_{AKGDH\_m}}{J_{PDH\_m}}$ = 0.177, $\mu$ = 0.22, $f_R$ = 0.037).

**Model Output**

*Aerobic and anaerobic growth rates as a function of $f_G$ and $f_R$*

$$\mu_{aero} = \mu(f_G, f_R, J_R > 0)$$

$$\mu_{anae} = \mu(f_G, f_R, J_R = 0)$$

The resulting $\mu$ surface is shown in (Fig. N4), with the global optima ($\mu_{aero}^{**}$ and $\mu_{anae}^{**}$) highlighted.

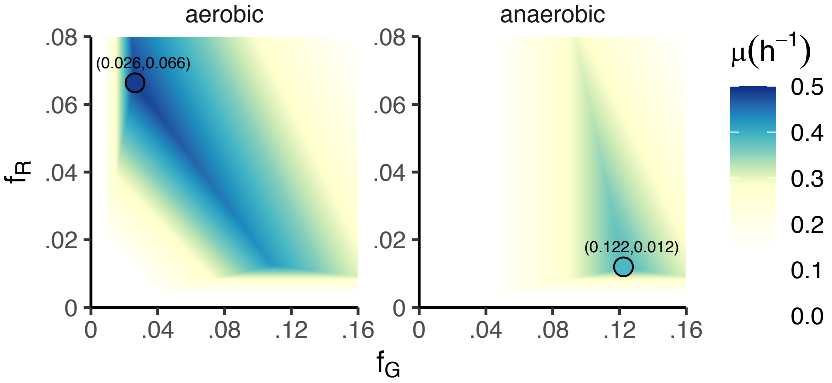

**Figure N4**| Aerobic or anaerobic growth rate $\mu$ as a function of the glycolytic ($f_G$) and respiratory protein abundance ($f_R$). Circles correspond to proteome compositions where $\mu_{aero}^{**}$ and $\mu_{anae}^{**}$ are achieved.

*Finding optimal aerobic growth rate and proteome allocation for a given $f_G$*

To accommodate potential future hypoxia, the glycolytic proteome may be expressed more than what is needed for optimal respiratory growth. Under the pressure of proteome hedging for oxygen depletion, yeasts may want to keep glycolytic proteome expression level at a certain value $f_G$. Under this constraint, optimal growth rate $\mu_{aero}^{*}(f_G)$

$$\mu_{aero}^{*}(f_G) = \mathbf{max}\ \mu_{aero}(f_G = f_G, f_R, J_R > 0)$$

$\mu_{aero}^{*}(f_G)$ is achieved only when [E10] and [E11] are both met, which means

$$\mu_{aero}^{*} = y_{ATP} * (2 * (J_G - s * \mu_{aero}^{*}) + \text{PO} * J_R) = E_T * (f_0 - f_G - f_R)$$

$$\mu_{aero}^{*} = \frac{2 * e_G * f_G + PO * e_R * f_R}{2s + 1/y_{ATP}} = E_T * (f_0 - f_G - f_R)$$

Define $e_T \equiv (2s + 1/y_{ATP}) * E_T$, thus the optimal growth rate is achieved with the optimal respiratory proteome

$$f_{R,aero}^{*} = \frac{e_T * f_0 - (e_T + 2 * e_G) * f_G}{PO * e_R + e_T} \qquad \text{[E13]}$$

And the optimal growth rate

$$\mu_{aero}^{*}(f_G) = \frac{E_T}{PO * e_R + e_T}[f_0 * PO * e_R + (2 * e_G - PO * e_R) * f_G] \qquad \text{[E14]}$$

*Finding optimal aerobic growth rate and proteome allocation*

$$\mu_{aero}^{**} = \mathbf{max}\ \mu_{aero}^{*}(f_G)$$

When $2 * e_G > PO * e_R$, i.e. glycolysis is more proteome efficient than respiration in ATP production, optimal growth is achieved with maximal $f_G$ (constrained by [E12]). When $PO * e_R > 2 * e_G$, i.e. respiration is more proteome efficient in ATP production than glycolysis, optimal growth is achieved with maximal $f_R$ (constrained by $J_{ETOH} = 0$).

*Finding optimal anaerobic growth rate and proteome allocation*

Optimal anaerobic growth rate $\mu_{anae}^{**}$ is achieved only when [E10] [E11] and [E12] are all met, which means

$$\mu_{anae}^{**} = y_{ATP} * 2 * (e_G * f_G - s * \mu_{anae}^{**}) = E_T * (f_0 - f_G - f_R) = f_R/\beta$$

thus, the glycolytic proteome fraction to achieve optimal anaerobic growth, $f_{G,anae}^{**}$, can be determined as

$$f_{G,anae}^{**} = \frac{e_T * f_0}{2 * e_G * (1 + \beta * E_T) + e_T} \qquad \text{[E15]}$$

*Predicting ethanol overflow from reserve glycolytic proteome capacity*

A key prediction of proteome-constrained model is the switch from full respiration to ethanol overflow when growth accelerates (Fig. 4f). This can be viewed as a growth optimization problem under the constraint of glucose uptake rate. To show that ethanol overflow emerges from reserve glycolytic proteome capacity, we add additional constraint that part of the anaerobic glycolytic proteome is retained. Define retained fraction of anaerobic glycolytic protein $r_G = f_G/f_{G,anae}^{**}$, and the growth optimization becomes

$$\mu(r_G, J_G) = \mathbf{max}\, \mu\left(f_G = r_G * f_{G,anae}^{**}, f_R, J_G = J_G, J_G < e_G * f_G\right)$$

Plots of $J_G \sim \mu$ and $J_{ETOH} \sim \mu$ at different $r_G$ show that, even though respiration is intrinsically more proteome efficient than glycolysis, when there is a need to conserve glycolytic proteome, optimal growth is achieved with overflow metabolism (full utilization of that glycolytic capacity) (Fig. N5).

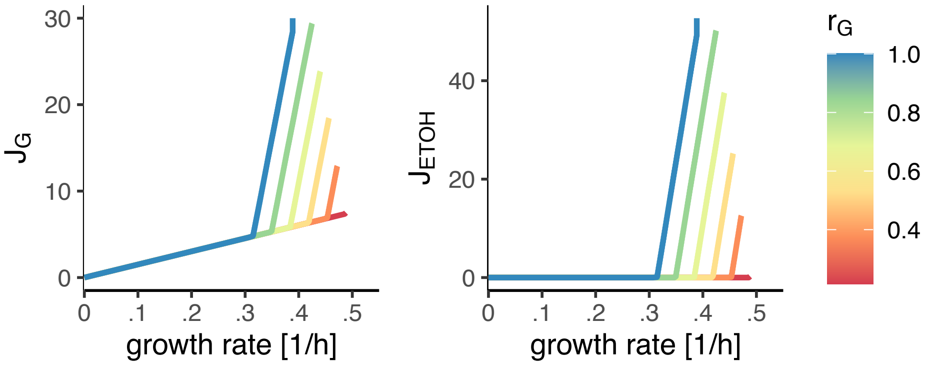

**Figure N5|** Predicted overflow as a function of varying retained glycolytic proteome ($r_G = f_G/f^{**}_{G,anae}$). $J_G$, glucose uptake rate [mmol/h/gDW]. $J_{ETOH}$, ethanol production rate [mmol/h/gDW].

**PO Ratio**

The amount of ATP produced per oxygen by respiration has an important impact on the efficiency of respiration versus glycolysis. For simplicity we used PO ratio of *S. cerevisiae* based on mechanistic efficiency of ATP synthase (3 ATP per 10 protons translocated across membrane, and 4, 2, 4 protons per pair of electrons pumped through complex I, III, IV, respectively; note that *S. cerevisiae* lacks complex I, and thus each O → 6 protons → 1.8 ATP). As mentioned in 'Finding optimal aerobic growth rate and proteome allocation', respiration is the optimal aerobic growth strategy as long as $PO * e_R > 2 * e_G$. With the $e_R$ and $e_G$ determined in this study (independent of PO), this means PO > 0.94.

One earlier study reported PO ratio = 0.95 in *S. cerevisiae*, which is close to this limit [2]. A higher PO ratio (PO = 1.4) has also been reported in *S. cerevisiae* [3]. As the lower PO ratio is widely used in many genome-scale models, we would like to revisit how it was obtained [2], which involved comparing growth-required ATP with oxygen consumption rate. The uncertain part is the ATP requirement. The uncertain part is the ATP requirement, which was determined based on a single experiment measuring glycolytic ATP production (based on glucose uptake rate) in an anaerobic, glucose-limited chemostat, without detailed methods provided, confidence limits, or evidence that this slow growing anaerobic condition generalizes. Indeed, the authors themselves noted evidence for a higher PO ratio: '…a P/O-ratio of 1.8-2.0 leads to a $Y_{ATP}$ of approximately 10-11 g biomass/mol ATP, which is generally believed to be the correct value for anaerobic growth of yeasts on glucose.'

We sought also to quantify the PO ratio, without any assumption on GAM or the mechanistic efficiency of the ETC or ATP synthase. To do that, we used glucose consumption rate and oxygen consumption rate (in batch culture) and PO ratio (the parameter of interest) to obtain total ATP production rate ($J_{ATP}$) and its standard error ($\sigma_{ATP}$) according to the stoichiometric flux model, and then obtained GAM by non-negative linear fitting of $J_{ATP}$ to growth rate $\mu$ ($\widetilde{J_{ATP}} = \text{GAM} * \mu + \text{NGAM}$, where NGAM is the non-growth associated ATP maintenance and $\widetilde{J_{ATP}}$ is the fitted value) across several conditions. We then used a Bayesian approach to obtain posterior PO distribution by calculating likelihood $\mathcal{L}(\text{PO}) = \prod_k p(\widetilde{J_{ATP}}(\text{PO}) \sim \text{norm}(J_{ATP}, \sigma_{ATP}))$. To improve confidence, we included conditions with varying levels of respiration: batch cultured *S. cerevisiae* CEN.PK (more respiratory) and FY4 (less respiratory), carbon-limited continuous culture (fully respiratory), and respiration-limited culture ($N_2$ sparged or antimycin treated, no respiration). With this, we obtained for *S. cerevisiae*, PO = 1.69, 95% confidence interval [1.590, 1.765].

For organisms whose mitochondria contain complex I (including *I. orientalis* and mammals), the PO ratio is higher (reported to be between 1.4 [2] and 2.5 [4]). Using similar approach, we estimated the PO ratio for *I. orientalis* as 2.15, 95% confidence interval [1.87, 2.39].
