## Supplementary material for "Proteome capacity constraints favor respiratory ATP generation": method

### Method for

#### Strains and Cultivation

*Strains*

The *I. orientalis* strain used in this study, SD108, is originally isolated from rotting bagasse[1]. Two prototrophic *S. cerevisiae* strains were used: CEN.PK2 (MATa) was a gift from Dr. Jose Avalos, Princeton, NJ, USA; and FY4 (DBY11069,MATa) that was derived from S288C background [2]. S288C strain naturally carries mutations that affect the gene HAP1 involved in respiratory regulation [3]. *I. orientalis* Δnde1 (Δg1781) and complex 1 mutant (Δg1702) was created with CRISPR/Cas9 editing [4,5] as described in reference [6].

*Media*

If not specified, yeasts were cultured in minimal media containing 20g/L glucose 6.7g/L yeast nitrogen base (YNB) without amino acids (pH = 5) (Sigma, Y0626). For nutrient limitation, YNB without amino acid or ammonium or phosphate (MP Biomedicals, 114029622) was used as the mineral base and supplemented with nutrient specified in Table below (adapted from reference [7]). All media were filter sterilized through a 0.22μm pore filter.

Table 1. culture media

| **media (g/L)** | **C limit** | **N limit** | **P limit** |
|---|---|---|---|
| YNB (-) AA (-) $NH_4$ (-) $PO_4$ | 0.700 | 0.700 | 0.700 |
| $(NH_4)_2SO_4$ | 5.000 | 0.050 | 5.000 |
| $KH_2PO_4$ | 1.000 | 1.000 | 0.010 |
| KCl | 0.000 | 0.000 | 0.542 |
| glucose | 0.800 | 5.000 | 5.000 |

*Measuring cell density by OD600*

To measure cell density, the culture was diluted 10 times with water and absorption at 600nm was measured by a UV-VIS spectrophotometer (GENESYS 10, Thermo). The measured absorption was then multiplied by 10 to obtain OD600. This ensures that the absorbance is kept within linear range.

*Culture*

For batch culture, the yeasts were first cultured overnight in minimal media to achieve final OD600 about 4 for *I. orientalis* and 3 for *S. cerevisiae*. For $^{13}C$ isotope tracing, the yeast was adapted in the same $^{13}C$ culture overnight to ensure isotopic steady state in the biomass. The overnight culture was then inoculated into 4mL media in 14mL round-bottom Falcon culture tube tilted at 45° angle or 20~40mL media in 150mL vented baffled culture flask at initial OD about 0.05~0.2, and cultured in a shaker at 250rpm and 30°C. Pseudo steady state is usually maintained below OD600 = 1.5.

For nutrient limited continuous culture, *S. cerevisiae* FY4 or *I. orientalis* SD108 was cultured in a home-built miniaturized multichannel bioreactor with a working volume of 20mL following the previous procedure [8]. 200μL overnight culture was inoculated in the culture tube and allowed to grow overnight before starting the continuous flow of media. The flow rate is controlled by a multichannel peristaltic pump (205S/CA12, Watson-Marlow, MA, US) and manifold tubing with proper internal diameter. The flow rate was calibrated each time by monitoring the effluent, and the volume of the culture was adjusted within ± 2mL to match the desired dilution rate. The cultures were mixed by sparging with 7.5 standard liters per min of water-saturated air for aerobic culture. The culture was maintained under continuous flow for at least 48hrs to achieve steady state. The final pH was measured to be about 3.5. Four dilution rates (0.08, 0.16, 0.22, 0.28 $h^{-1}$) were used for *S. cerevisiae* and five (0.12, 0.18, 0.23, 0.34, 0.45$h^{-1}$) for *I. orientalis* for each nutrient limitation.

Oxygen-depleted batch culture was done in the same home-build bioreactor, but without media feeding and with continuous sparging of 7.5 standard liters per min of water-saturated nitrogen.

For antimycin treatment, a concentrated stock of antimycin (100mM in DMSO) was first diluted 100X with water and then added to the culture at 100X dilution.

**Metabolite extraction**

Yeast metabolites were extracted by chilled solvent following rapid vacuum filtration. Specifically, a total amount of cell culture equivalent to 2mL at OD600 = 1 was extracted. Batch cultures were extracted at OD600 between 0.6 and 0.9. The cell culture was rapidly filtered through a nylon membrane filter (0.5µm pore size) on a fritted glass support of vacuum filter flask. Metabolism was quickly quenched by immersing the membrane in 1.5mL 40:40:20 acetonitrile:methanol:water with 0.5% formic acid prechilled in -20˚C in a 60mm petri dish. Extraction was allowed to continue on ice for 1min and then neutralized by 132 µL 15.8 g/L $NH_4HCO_3$ solution. The extract was used to rinse the membrane, transferred to a 1.5mL tube, and stored in -80˚C. The extract was centrifuged at 17,000g at 4˚C for 10min to obtain the supernatant for LC-MS analysis. For each growth condition, three extractions were obtained and analyzed.

**Determining major fluxes**

*Determining metabolite concentrations in culture supernatant*

Glucose, ethanol, acetate, succinate, and glycerol in spent media were measured with $^1$H-NMR (500 MHz Advance III, Bruker, MA, US). 50mM TMSP-$d_4$ internal standard was 1:10 diluted into the spent media for internal reference. A fresh media sample was also included to calibrate TMSP. $^1$H-NMR spectra were collected using the following acquisition parameters: TD = 65536, NS = 64, D1 = 5s, O1P = 4.68, P1 = 11.69, P12 = 2400, SPW1 = 0.002, SPNAM1 = Gaus1_180r.1000. Chemical shift used for quantification: 0 ppm (s, 9H) for the TMSP standard, 3.22 ppm (dd, 1H) for glucose, and 1.17 ppm (t, 3H) for ethanol, 2.07ppm (s, 3H) for acetate, 2.60 (s, 4H) for succinic acid, and 3.64 (m, 4H) for glycerol. Quantitation was done in MestReNova.

Glucose concentration was also determined by a biochemistry analyzer (2900, YSI, OH, US). Spent media with initial glucose concentration of 20g/L was measured with 4-fold dilution to be within linear range. Each sample was measured with at least two technical replicates.

*Growth rate and extracellular fluxes for pseudo-steady-state batch culture*

Growth rate ($\mu$) and metabolite fluxes (j) in batch culture were determined by sampling the culture at least four times (t) during the exponential growth phase, starting from OD600 about 0.1 after allowing the culture to adapt in the fresh media (about 1 hrs in aerobic culture and 4hrs in anaerobic or antimycin treatment), to OD600 about 1.5 or before half of the glucose was consumed. At each time point, OD600 was measured and supernatant was saved for analysis of metabolite concentration (c). Growth rate ($\mu$) was determined with linear fitting: $\mu$ = slope (ln OD ~ t), while extracellular flux (j) was the product of growth rate and the slope of c ~ OD, j = $\mu$ * slope (c ~ OD). The resulting j is in unit of mmol/L/OD600/h, which is then converted to mmol/gDW/h using the OD-to-biomass conversion factor determined in biomass analysis (e.g. for batch culture, this conversion factor is around 0.35 gDW/L/OD600 for both yeasts). Errors are determined by propagating error from the linear regression.

*Oxygen consumption rate*

Oxygen consumption rate was measured for batch culture with a Clark-type dissolved oxygen probe (B40PCID, 89231-624, VWR, PA, US). The culture was kept in a glass chamber and temperature was maintained by water bath. The culture was fully oxygenated first, and then sealed with the temperature-equilibrated probe. Dissolved oxygen was measured every 20sec for 5 min or until oxygen drops to 60% saturation. During the measurement, the culture was gently mixed by magnetic stirrer. The culture density used for measurement was OD600 = 0.2 ~ 0.3 for *I. orientalis* and OD600 = 0.6 ~ 0.8 for *S. cerevisiae*. Oxygen consumption rate was then calculated by linear fitting of the oxygen concentration change over time, and normalized by the cell density.

*Flux determination in continuous culture*

After steady state was reached, the continuous culture was sampled by collecting 1mL effluent at least three times over 12hr. For each sampling, OD600 was measured and remaining glucose concentration was determined by YSI biochemistry analyzer. The whole culture was then cooled on ice and centrifuged in 4°C. The metabolite concentration (c) was determined in the supernatant. Fluxes (j) were then calculated as $j = dr * (c_0 - c)$, where dr is the dilution rate, $c_0$ is the initial concentration in the media, and then normalized by biomass concentration.

**Dry weight and biomass composition**

*Calibrating biomass composition in a reference yeast culture*

We first measured DNA, RNA, protein, and carbohydrate in a reference condition, *S. cerevisiae* in carbon limitation at $0.1h^{-1}$, using method described previously [6]. Briefly, protein content was determined using the Biuret method with BSA (23209, Thermo Fisher, MA, US) calibration. Cell pellet equivalent to 1mL of OD600 = 1 was washed and lysed in 300μL 1M NaOH at 98 °C for 5min. 100μL 1.6% $CuSO_4$ was then added to the lysate and absorbance at 555nm was used to quantify protein concentration. For RNA quantitation, the cell pellet was lysed in 300μL 0.3M KOH at 37°C for 60 min, and then 100μL 3M $HClO_4$ was added to precipitate DNA and protein. The supernatant was combined with 600uL 0.5M $HClO_4$ used to wash the precipitant, and RNA content was then determined by the absorption at 260nm with pathlength correction (1cm), using an extinction coefficient of 31 μg/mL/A260. For DNA, cell pellet equivalent to 10mL of OD600 = 1 was hydrolyzed with 500μL 1.6 M $HClO_4$ for 30 min at 70˚C, and allowed to react with 1 mL diphenylamine reagent (0.5 g diphenylamine in 50 mL acetic acid, 0.5 ml 98% $H_2SO_4$, and 0.125 ml 3.2% acetaldehyde water solution) at 50˚C for at least 3 hours. Absorption at 600nm was measured from the supernatant, and used to quantify DNA concentration with the calibration of a purified DNA standard (15633019, ThermoFisher Scientific, MA, US).

*LC-MS assay for measuring biomass*

Protein, DNA, RNA and carbohydrate in biomass is measured by acidic hydrolysis with reference to $^{13}C$ labeled *S. cerevisiae*, which is cultured in carbon limitation with [U-$^{13}C_6$] glucose at $0.1h^{-1}$. To determine biomass in a given sample, three replicates of 1mL culture was pelleted and each combined with an aliquot of $^{13}C$ *S. cerevisiae* equivalent to 1mL of OD600 = 1. The pellet was washed with water twice, and hydrolyzed in 100μL 6M HCl at 80 °C and 300 rpm for 2h with a thermomixture. The hydrolysate was then centrifuged, and 8μL of supernatant was dried with nitrogen gas, and re-dissolved in 80μL 40:40:20 acetonitrile:methanol:water for LC-MS analysis. The detected monomers were categorized into components of protein, DNA, RNA, and carbohydrate (Supplementary Table), and the $^{12}C/^{13}C$ ratio from each category was averaged to obtain concentration relative to the reference condition.

Lipid in biomass is analyzed by saponification and quantified by spiking in a mixture of $^{13}C$ fatty acids of highest abundance in yeast. Specifically, 3 replicates of 1mL culture was pelleted, and saponified in 1mL 0.3M KOH in 10:90 water:methanol containing internal $^{13}C$ standard of 40μM [U-$^{13}C_{16}$] palmitate, 40μM [U-$^{13}C_{18}$] oleate, and 20μM [U-$^{13}C_{18}$] linoleate for 1h at 80 °C. The mixture was then acidified by 100μL formic acid and extracted twice with 1mL hexane. The upper layer was separated, dried under nitrogen gas, redissolved in 100μL 1:1 acetonitrile:methanol, and analyzed by reverse phase LC-MS. Fatty acids were quantified by $^{12}C/^{13}C$ ratio for the three with internal reference, and by MS peak intensity for other fatty acid species.

**Metabolite analysis by LC-MS**

*Metabolite extraction*

For extracting intracellular metabolites, yeast culture equivalent to 3mL of OD600=0.8 was vacuum filtered using Nylon membrane filters (0.5μm pore size, 1213776, GVS Magna™), and then quickly immersed in 1.5mL 40:40:20 acetonitrile:methanol:water with 0.5% formic acid precooled in -20°C. After about 1min incubation on ice, the extract was neutralized by 132uL 15.8% $NH_4HCO_3$. The extract was then centrifuged at 17000rpm at 4°C to obtain supernatant ready for LC-MS analysis for polar metabolites. For each sample, three independent extractions were made.

*HILIC LC-MS for polar metabolites*

Separation of polar metabolites was achieved with hydrophilic interaction chromatography (HILIC), using a Vanquish UHPLC system (Thermo Fisher Scientific, CA, US) and an XBridge BEH Amide column (2.1 mm x 150 mm, 2.5 mm particle size, 130 A˚pore size; Waters, Milford, MA). The LC runs at flow rate of 150μL/min with a 25 min solvent gradient as following: 0 min, 85% B; 2 min, 85% B; 3 min, 80% B; 5 min, 80% B; 6 min, 75% B; 7 min, 75% B; 8 min, 70% B; 9 min, 70% B; 10 min, 50% B; 12 min, 50% B; 13 min, 25% B; 16 min, 25% B; 18 min, 0% B; 23 min, 0% B; 24 min, 85% B; 30 min, 85% B, where solvent A is 95:5 water:acetonitrile with 20 mM ammonium hydroxide and 20 mM ammonium acetate, pH 9.4, and solvent B is acetonitrile. Autosampler temperature was 4ºC, column temperature was 25 ºC, and injection volume was 10μL. LC was coupled to a quadrupole-orbitrap mass spectrometer (Q Exactive, Thermo Fisher Scientific, CA, US) via electrospray ionization. The mass spectrometer operates in negative and positive ion switching mode and scans from m/z 70 to 1000 at 1 Hz and 140,000 resolution, with additional selected ion monitoring scan from m/z 650 to 770 for NAD(P) cofactors. Data was collected by XCalibur (Thermo Fisher Scientific).

*Reverse-phase LC-MS for saponified fatty acid*

Saponified fatty acids were analyzed by liquid chromatography (Accela U-HPLC) coupled with orbitrap mass spectrometer (Exactive, Thermo Fisher Scientific, CA, US). LC separation was done by reverse-phase ion-pairing through a Luna C8 column (150 × 2.0 $mm^2$, 3 μM particle size, 100 Å pore size; Phenomenex) with a solvent gradient of 0 min 80% B; 10 min, 90% B; 11 min, 99% B; 25 min, 99% B; 26 min, 80% B; 30 min, 80% B, where solvent A is 10 mM tributylamine + 15 mM acetic acid in 97:3 H2O:methanol, pH 4.5, and solvent B is methanol. The flow rate was 250 μL/min and column temperature 25 °C with an injection volume of 5 μL. The MS scans were in negative-ion mode with a resolution of 100,000 and scan range of m/z 120–600. Data was collected by XCalibur (Thermo Fisher Scientific).

*Metabolite quantitation from LC-MS*

Raw LC-MS data were converted to mzxml format by ProteoWizard (https://proteowizard.sourceforge.io). Peak picking and quantitation were done in the El-Maven software (v.0.4.1, Elucidata). For comparing across nutrient conditions, samples were extracted and analyzed in the same day to reduce batch effect. Relative fold change of each metabolite was quantified by relative peak area top in the chromatogram. For comparing between different yeasts in batch culture, $^{12}C$ labeled *S. cerevisiae* was 1:1 mixed with $^{13}C$ labeled *I. orientalis*, and $^{13}C$ labeled *S. cerevisiae* was 1:1 mixed with $^{12}C$ labeled *I. orientalis*. For each compound, ratio between labeled and unlabeled peaks was used for relative quantitation. For samples with $^{13}C$ labeling, natural isotope abundance was corrected using AccuCor [9] (https://github.com/lparsons/accucor).

#### $^{13}$C isotope tracing and genome-scale $^{13}$C metabolic flux analysis

##### *$^{13}$C isotope tracing*

Two glucose tracers were used for flux analysis: [U-$^{13}C_6$] glucose and [1,2-$^{13}C_2$] glucose, each was mixed with unlabeled glucose to achieve 1:1 molar ratio (50% enrichment). Batch cultures were adapted in the tracer media overnight before allowing to grow in fresh media for more than 3 generations. Continuous cultures were cultured in tracer media for the whole experimental period. 13C mass isotopomer distribution in about 40 metabolites was then analyzed by LC-MS from extracted polar metabolites and biomass hydrolysates (see *LC-MS assay for measuring biomass*).

##### *Genome-scale carbon mapping model for $^{13}$C-MFA*

We developed new carbon mapping models for *S. cerevisiae* and *I. orientalis* based on their genome-scale models, *iIsor*850 for *I. orientalis* [6] and *iSace*1144 for *S. cerevisiae* (reformatted from yeast 8.3.4 model [10] as described previously [11], available at https://github.com/maranasgroup/iSace_GSM). For model reduction, flux variability analysis [12] was performed with constraints on measured glucose uptake and byproduct (ethanol, acetate, glycerol, and succinate) excretions, to remove reactions incapable of carrying flux under glucose utilizing conditions, e.g. degradation pathways that form ATP-consuming futile cycles with biosynthesis of nucleotide, lipid, fatty acid, and carbohydrate. We also simplified intracellular compartments by assigning non-mitochondrial reactions to cytosol. Carbon mapping of reactions were obtained from previous large-scale mapping model in E.coli [13], or for new reactions curated from BioCyc database [14], biochemistry textbook, and the literature. Annotation of functional groups and adjacent carbon atoms were also provided for carbon atoms (which were previously associated with only numbering indexes) to facilitate future use. The models also contain cofactor balance (e.g., ATP, NADH), as well as proton pumping and the electron transport chain pathway. Stoichiometry of the 52 precursors in the biomass reactions are updated to reflect condition-specific macromolecular composition measured in this study. The mapping model for *S. cerevisiae* contains 394 reactions and 354 metabolites, while the model for *I. orientalis* contains 386 reactions and 363 metabolites. Both models, additional annotation, and biomass worksheet are available at https://github.com/maranasgroup/yeastsMFA

##### *$^{13}$C-metabolic flux analysis*

We employed $^{13}C$-MFA procedure described previously [13] (formulated using the elementary metabolite unit framework [15], available at https://github.com/maranasgroup/SteadyState-MFA). Briefly, a non-linear optimization formulation was used to find a flux solution by minimizing the sum of squared differences between the simulated $^{13}C$ mass isotopomer distributions (as a function of fluxes) and the observed ones from both tracers, as well as uptake/excretion fluxes. The best-fit flux solution was chosen from 100 randomized initialization. Goodness-of-fit test (chi-square) and 95% confidence interval estimation were performed as described previously [16,17]. All the described input data and MFA evaluation are available at https://github.com/maranasgroup/yeastsMFA.

#### Quantitative proteomics

Absolute protein abundance in batch cultured yeasts was quantified by label-free intensity-based absolute quantitation (IBAQ) using a UPS2 internal standard. Relative protein abundance was quantified across nutrient conditions or following antimycin treatment using TMTpro isobaric tags [18].

*Proteomics sample preparation*

Proteomics samples were prepared as previously described with modification [19,20]. Yeast pellet equivalent to 20mL OD600 = 1 was grounded by a cryomill (Retsch, Newtown, PA) at 25Hz for 10min and lysed in 50 mM HEPES pH 7.2, 4% SDS, and 1 mM DTT. Protein concentrations were determined in the supernatant of the lysate by BCA assay (Pierce BCA Protein Assay Kit, Thermo Scientific), and 300 µg protein were aliquoted and spiked with 2.5 µg of UPS2 Dynamic Range Standard (Sigma). The sample was then reduced with 5 mM dithiothreitol for 20 min at 60 °C and alkylated with 20 mM N-ethylmaleimide for 20 min at room temperature. 5 mM dithiothreitol was added to quench the excessive alkylating reagents. Proteins were purified by methanol-chloroform precipitation. The dried pellet was resuspended in 10 mM EPPS (*N*-(2-Hydroxyethyl)piperazine-*N*'-(3-propanesulfonic acid, pH 8.5) with 6 M guanidine hydrochloride). Samples were heated at 60 °C for 15 minutes, and the protein mixture was diluted 3-fold with 10 mM EPPS (pH 8.5). The protein mixture was digested with 6 µg LysC (Wako) overnight at room temperature. Samples were further diluted 4-fold with 10 mM EPPS (pH 8.5) and digested with an additional 20 ng/µL LysC and 10 ng/µL sequencing grade Trypsin (Promega) at 37 °C for 16 hours. Supernatant that contains peptides was obtained by ultracentrifugation at 100,000 rcf for 1 hour at 4 °C (Beckman Coulter, 343775), and then vacuum-dried. The dried peptides were resuspended and desalted using homemade stage tips with C18 material (Empore). The samples were resuspended in 1% formic acid to 1 µg/µL before LC-MS analysis.

For TMT labeling, pre-mixed TMTpro tags (16-plex and 18-plex, Thermo Scientific, 20 µg/µL in dry acetonitrile stored at -80 °C) were added at 5µg TMTpro : 1µg peptide ratio to the above supernatant containing 200 µg of peptides, mixed, and incubated at room temperature for 2 hours. The reaction was then quenched by addition of 5 µL of 5% hydroxylamine (Sigma, HPLC grade) at room temperature for 30 minutes. The resulting mixture was vacuum-dried, desalted, and resuspended as described above for LC-MS analysis.

For label-free quantification, one replicate of each yeast strain was also analyzed after prefractionation to detect a larger number of peptides. Specifically, prior to LC-MS analysis, the dried peptides were resuspended in 10 mM ammonium bicarbonate (pH 8) with 5% acetonitrile to a peptide concentration of 1 µg/uL. The dissolved peptides were separated into 96 fractions using a medium pH reverse phase separation (Zorbax 300Extend C18, 4.6 x 250 mm column) on a 1260 Infinity II LC system (Agilent) as described previously [20]. Each resulting 96-well plate was combined into 24 fractions [21], and each fraction was desalted and resuspended for LC-MS analysis as above.

*Peptide analysis by LC-MS*

Samples were analyzed on an EASY-nLC 1200 HPLC (Thermo Fisher Scientific) coupled to an Orbitrap Fusion Lumos mass spectrometer (Thermo Fisher Scientific) with Tune version 3.3. Data was collected by XCalibur (Thermo Fisher Scientific). Peptides were separated on an Aurora Series emitter column (25 cm × 75 µm ID, 1.6 µm C18) (Ionopticks, Australia) and held at 60°C using an in-house built column oven. Solvent A consisted of 2% DMSO (LC-MS grade, Life Technologies), 0.125% formic acid (98%+, TCI America) in water (LC-MS grade, OmniSolv, VWR), solvent B of 80% acetonitrile (LC-MS grade,

OmniSolv, Millipore Sigma), 2% DMSO and 0.125% formic acid in water. The following 90 min-gradient was applied at a constant flow rate of 350 nL/min after thorough equilibration of the column to 0% B: 0% – 6%B in 5 min; 6 – 25%B for 70 min; 25% – 100% for 10 min; 100% for 5 min.

For electrospray ionization, 2.6 kV were applied between 1min and 83min of the LC gradient. The Fusion Lumos was operated in data dependent mode. The survey scan was performed at a resolution setting of 120k in orbitrap, followed by MS2 duty cycle of 1.5 s. The normalized collision energy for CID MS2 experiments was set to 30%, and the HCD collision energy was set at 24%. The ion trap detector was used for MS2 scans. To avoid carry-over of peptides, 2,2,2-trifluoroethanol (>99% Reagent plus, Millipore Sigma) was injected in a 30 min wash between each sample.

*Proteomics data analysis*

The data was analyzed using GFY software licensed from Harvard University. Raw files were converted to mzXML using ReAdW.exe. MS2 spectra assignment was performed using the SEQUEST algorithm v.28 (rev. 12) by searching the data against the combined reference proteomes for *S. cerevisiae* (S288C: UP000002311, 2/24/2021; CEN.PK, UP000013192, 8/20/2021) and *I. orientalis* (UP000029867, 11/13/2019) acquired from Uniprot merged with the UPS2 Proteomics Standards FASTA file (https://www.sigmaaldrich.com/deepweb/assets/sigmaaldrich/marketing/global/fasta-files/ups1-ups2-sequences.fasta) along with common contaminants such as human keratins and trypsin. The target-decoy strategy was used to estimate the peptide false discovery rate (FDR)[22], and 1% FDR cutoff was used for MS2 spectral assignment. A 20-ppm precursor ion tolerance with the requirement that both N- and C-terminal peptide ends are consistent with the protease specificities of LysC and Trypsin was used for SEQUEST searches. One missed cleavage was allowed, and NEM was set as a static modification of cysteine residues (+125.047679 Da). Fragment ion tolerance in the MS2 spectrum was set at 1 Th. Filtering was performed using a linear discriminant analysis with the following features: Sequest parameters XCorr and unique ΔXCorr, peptide length, missed cleavages, adjusted PPM, fraction of ions matched, and charge state. Forward peptides within three standard deviations of the theoretical m/z of the precursor were used as positive training set. All reverse peptides were used as negative training set. Linear Discriminant scores were used to sort peptides with at least seven residues and to filter with the desired cutoff. Furthermore, we performed a filtering step on the protein level by the "picked" protein FDR approach[23]. Protein redundancy was removed by assigning peptides to the minimal number of proteins which can explain all observed peptide, with above-described filtering criteria [24].

Relative quantification of TMT tagged samples was done by summing up area of TMT reporter ion belonging to each protein. The signal was then normalized to the mean across samples, and then median normalized within each sample. To quantify absolute protein abundances in label-free samples, for each protein, area of precursor ion intensity from all peptides was summed up, and then normalized by number of theoretical peptides. Signals from UPS2 proteins were used to construct a calibration curve, which was then fitted to a power law, log(intensity) = k * log(concentration) + constant, to obtain absolute concentration of yeast proteins (log-linear coefficient $k = 1.25 \pm 0.08$ on average). Absolute protein abundance in batch culture is reported as mass fraction in whole proteome, which is approximated by the product of concentration and amino acid sequence length normalized to the sum of all proteins. Absolute protein abundance in nutrient limitation or respiratory deficient conditions is inferred from the relative fold change to batch culture obtained with relative quantification.

**Multi-omics integration and identification of metabolic regulators in *I. orientalis***

*Systematic identification of meaningful metabolic enzyme regulation (SIMMER)*

Integration of fluxomics, metabolomics, and proteomics data were performed as described previously with adaptation for *I. orientalis* [25]. A total of 241 metabolites were measured, 129 of which were matched to genome-scale model by ChEBI compound identifier. A total of 1922 proteins were quantified, 418 of which were enzymes in the genome-scale model according to gene-protein-reaction (gpr) annotation. Flux through 343 reactions were obtained from $^{13}$C MFA. For 51 of these reactions, omics data were used to fit rate laws and identify regulation. These reactions have confidently determined flux (median of (confidence interval / best fit) across all conditions < 1), no more than 1 organic reactants (substrates or products) unmeasured, at least 1 substrate measured, and at least 1 gene measured in proteomics. For each reaction, kinetic models with or without putative regulator were generated based on Michaelis-Menten kinetics as described previously [25,26]. These putative regulators were drawn from BRENDA database [27], and include measured metabolites not necessarily in the metabolic model. For each reaction, parameters ($k_{cat}$, $K_d$, $K_{eq}$) were inferred by fitting flux, metabolite and enzyme concentrations across conditions with different kinetic models, using a non-negative least square Monte-Carlo Markov chain (NNLS-MCMC) procedure described previously (code available at https://github.com/shackett/simmer) [25]. Choice of priors are same as described previously [25], and MCMC sampling is done every 20 samples for a total of 1000 samples desired with first 200 samples discarded. Regulators were identified at false discovery rate 0.1 with a likelihood ratio test between kinetic models with and without regulator, based on the expectation that increase in fit (2*(difference in log likelihood)) follows a chi-squared distribution with 1 degree of freedom. As biochemical analysis on *I. orientalis* enzymes were largely unavailable, unlike previous study in *S. cerevisiae* [25], no prior biochemical knowledge of yeast regulation was implemented.

*Metabolic leverage*

Metabolic leverage measures how much of flux variation across environmental conditions is accounted for by variation in individual species of a reaction [25]. Metabolic leverage of reaction species $k$ is calculated as

$$\psi_k = \frac{\left(\left.\frac{\partial v}{\partial s_k}\right|_{s_k=\overline{s_k}}\right)^2 \mathrm{Var}_e(s_k)}{\sum_{k=1}^{n}\left(\left.\frac{\partial v}{\partial s_k}\right|_{s_k=\overline{s_k}}\right)^2 \mathrm{Var}_e(s_k)}$$

where $\left.\frac{\partial v}{\partial s_k}\right|_{s_k=\overline{s_k}}$ is the sensitivity of reaction flux to component k at its mean concentration obtained from best supported kinetic model (and parameters), and $\mathrm{Var}_e(s_k)$ is the variance of species $k$ across environmental conditions.

**Validation of ATP inhibition of glyceraldehyde-3-phosphate dehydrogenase (GAPD)**

*Purification of I. orientalis GAPD*

The GAPD sequence of *I. orientalis* (XP_029320181.1, amino acid 2-335) was synthesized by Genscript as a codon optimized version for E. coli expression with an N-terminal HIS-Tag followed by an rTEV cleavage site and cloned into pET24. After transformation into E. coli B21[DE3], a 20 ml culture was induced at OD600 ~ 1.0 with 1 mM IPTG and grown further overnight at 30 °C. Protein purification was performed using small scale NI-NTA Spin Kit (Qiagen, #31314) using the manufacturer's protocol for native conditions to obtain purified GAPDH at about 1mg/mL (estimated by A280).

*GAPD enzyme activity assay*

GAPD enzyme activity was measured by NADH absorption through an arsenate coupled reaction adapted from an earlier study [28]. Specifically, reaction mixture contains 100mM Tris HCl (pH 8.5), 800uM EDTA,

25mM $Na_2HAsO_4$ (Sigma, S9663), and 1mM NAD+ (Sigma, N7004). 1mM GAP (Cayman, 17865) was used for calibrating enzyme activity while 150μM GAP for measuring ATP inhibition. The latter is approximately physiological concentration according to reported absolute concentration in *S. cerevisiae* [26]. Enzyme stock was first activated by adding 10mM dithiothreitol, and then diluted into reaction mixture to achieve working concentration at about 2U/L (about 50- to 200-fold dilution). To improve accuracy, GAP and enzyme was individually diluted to 2X working concentration in the reaction mixture, and the reaction was initiated by 1:1 mixing the 2X substrate mix and 2X enzyme mix. NADH absorption at 340nm was read every min by a multi-well plate reader (BioTek, Synergy HT). Initial reaction rate was obtained by linear fitting. For ATP inhibition, different concentration of ATP (Sigma, A6419) was added to the substrate mix before combining with enzyme mix.

**Proteome efficiency**

*ATP flux*

For yeasts, glycolytic ATP production is calculated as PGK_c + PYK_c − HEK_c − PFK_c flux. Respiratory ATP production is represented by the flux through ADPATPt_c_m, the mitochondrial ADP/ATP transporter. 95% confidence interval [lb, ub] is obtained from $^{13}C$ MFA, based on which the ATP flux is determined as (lb + ub)/2 with a standard error of (ub-lb)/3.84.

For NCI-60 cancer cell lines, the flux data was obtained from a previous flux analysis, using a model that assumes 4 proton translation per ATP production from the ATP synthase [29], and constrained by experimentally measured rates (growth, uptake and excretion) [30]. ATP production is obtained similar to yeast.

Flux in mouse tissues was obtained from a recent study that measured TCA flux and glucose uptake in vivo[31]. Respiratory ATP flux was calculated as 14.5 ATP per AcCoA oxidized in TCA cycle (as shown previously [31]), while glycolytic ATP flux was based on 2 ATP per glucose. Wet tissue mass is converted to dry mass by a factor of 0.4, and protein is assumed to account for 0.5 of dry mass.

*Pathway protein mass fraction*

Each *S. cerevisiae* protein is assigned to a functional sector based on gene ontology from Uniprot and pathway annotation from the genome-scale model. *I. orientalis* proteins were assigned based on protein sequence identity to *S. cerevisiae* obtained from blastp (https://blast.ncbi.nlm.nih.gov/Blast.cgi?PAGE=Proteins). Functional assignment in other organisms were obtained similar to *S. cerevisiae*. The complete list of functional assignment can be found in Supplementary Table. To obtain pathway proteome mass fraction in total cell dry weight, measured biomass composition was used for yeast. For organisms where biomass composition is not measured, protein is assumed to account for 50% of dry mass.

*Flux partitioned proteome allocation*

Since glycolysis is used to provide precursor for respiration, and both glycolysis and respiration are used to provide biomass precursors, proteome allocation required for 'fermentation' (converting glucose to ethanol) and 'respiration' (converting glucose to CO2) was calculated based on flux partitioning [34]. Briefly, the protein cost of enzyme $i$ , $f^i$, is divided among fermentation ($f$), respiration ($r$), biomass ($bm$). Its cost for function $k$, $f_k^i$, is proportional to the carbon flux its product is used for $k$, $j_k$

$$f_k^i = f^i \cdot \frac{j_k}{\sum_k j_k}$$

$j_r$ is approximated by oxygen consumption rate; $j_f$ is approximated by 3-times of ethanol excretion flux; $j_{bm}$ is derived from precursor stoichiometry in biomass equation. For simplicity, oxphos is not required for biomass precursors.

**Dissolved oxygen measurement in a plate culture**

To measure dissolved oxygen, exponential phase yeast culture was added to 100uL fresh media in a 96-well plate coated with phosphorescence oxygen sensor at the bottom (OxoPlate, OP96U, PreSens Precision Sensing GmbH, Germany). The culture was allowed to accommodate for 10min and then analyzed by a plate reader with or without fast shaking at 30 °C (BioTek, Synergy HT). Calibration and measurement were done following the manufacturer's procedure.

**Competitive co-culture and fitness**

*Co-culture and genomic DNA extraction*

Overnight cultures of *S. cerevisiae* CEN.PK and *I. orientalis* SD108 were mixed at 1:1 according to OD600, pelleted, and inoculated into fresh media at OD600 = 0.5. The cultures were then grown aerobically in one of the following conditions: aerobic 10 g/L ethanol, 20 g/L glucose, 20 g/L sucrose YNB with serial transfer for 12 ~ 14 hrs; or in aerobic glucose-, ammonia-, or phosphate-limited continuous culture at 0.1 h-1 dilution rate for 24 hrs. For (cyclically) anaerobic culture in glucose, anaerobic phase was achieved by sparging nitrogen into the culture at desired duty cycle (75%, 18h/24h; 87%, 21h/24; 100%, 24h/24h). Anaerobic culture was achieved by sparging nitrogen into the culture with 20g/L glucose. Relative abundance of the two yeasts was measured by qPCR at 4 to 6 time points and used to obtain fitness.

At each time point, 1mL co-cultures was pelleted, and the rest was then diluted with fresh media to keep the OD600 of the culture to approximate 1 in aerobic cultures or 0.5 in (cyclically) anaerobic cultures. For time points sampled, see Fig. S7. Calibration curve was also prepared by mixing single cultures at different ratios. The cell pellet was lysed by lyticase (Sigma Aldrich, L4025) and genomic DNA was extracted with DNeasy® Blood & Tissue Kit (Qiagen) following the manufacturer's procedure.

*Determine relative species abundance by quantitative PCR (qPCR)*

Relative abundance of *S. cerevisiae* and *I. orientalis* was determined by qPCR of the pho2 genomic sequences for *S. cerevisiae* (Gene ID: 851452, NM_001180165.1) and the distant homolog for *I. orientalis* (Gene ID: 40382003, XM_029463910). qPCR primers were designed using OligoArchitect Online (Sigma-Aldrich, http://www.oligoarchitect.com) and checked for cross-hybridization against the genomic sequences of both species using BLAST (https://blast.ncbi.nlm.nih.gov/Blast.cgi). The primers and probes used for *S. cerevisiae* are CTCTCTTCTTCGATCATG (sense), TCTCCTCATTATTAGCATTATG (anti-sense), and [6FAM]ATAACCAACACCAACAACGGACAAG[OQA] (probe); for *I. orientalis*, GAGACTAGCACCCTTAAC (sense), CGTTCACATCTACACTGA (anti-sense), and [JOE]ACAGCCTCCACAACGACTTCT[TAM] (probe) (Sigma-Aldrich). Primers and probes were tested for unspecific cross-reactivity using iTaq™ Universal Probes Supermix (BioRad 1725132) individually and in combination with genomic DNA from both species in various ratio. No cross-activity was observed. For qPCR, 1 to 2 ng of isolated DNA was used per 10 µL assay containing 250 nM for each primer and

125 nM each probe) in a 384 plate. Assays were performed using the Applied Biosystems ViiATM7 Real-Time PCR System. The relative abundance was quantified from the calibration curve, and fitted to log(I.o / S.c) = fitness * t + constant to obtain relative fitness.

**Data and code availability**

All source data are provided in Supplementary Table. Raw proteomics data will be made available in PRIDE (https://www.ebi.ac.uk/pride/).

Data analysis and visualization were done in R (version 3.5.1) and Matlab (version 2021b). R code for multi-omic integration and metabolic regulation analysis is available at https://github.com/yihuishen/yeast_regulation. Yeast metabolic models and code for MFA are available at https://github.com/maranasgroup/yeastsMFA and https://github.com/maranasgroup/SteadyState-MFA
